# Cdk1-dependent destabilization of long astral microtubules is required for spindle orientation

**DOI:** 10.1101/2020.05.23.111989

**Authors:** Divya Singh, Nadine Schmidt, Franziska Müller, Tanja Bange, Alexander W. Bird

## Abstract

The precise execution of mitotic spindle orientation in response to cell shape cues is important for tissue organization and development. The presence of astral microtubules extending from the centrosome towards the cell cortex is essential for this process, but little is understood about the contribution of astral microtubule dynamics to spindle positioning, or how astral microtubule dynamics are regulated spatiotemporally. The mitotic regulator Cdk1-CyclinB promotes destabilization of centrosomal microtubules and increased microtubule dynamics as cells transition from interphase to mitosis, but how Cdk1 activity specifically modulates astral microtubule stability, and whether it impacts spindle positioning, is unknown. Here we uncover a mechanism revealing that Cdk1 destabilizes astral microtubules to ensure spindle reorientation in response to cell shape. Phosphorylation of the EB1-dependent microtubule plus-end tracking protein GTSE1 by Cdk1 in early mitosis abolishes its interaction with EB1 and recruitment to microtubule plus-ends. Loss of Cdk1 activity, or mutation of phosphorylation sites in GTSE1, induces recruitment of GTSE1 to growing microtubule plus-ends in mitosis. This decreases the catastrophe frequency of astral microtubules, and causes an increase in the number of long astral microtubules reaching the cell cortex, which restrains the ability of cells to reorient spindles along the long cellular axis in early mitosis. Astral microtubules must thus not only be present, but also dynamic to allow the spindle to reorient in response to cell shape, a state achieved by selective destabilization of long astral microtubules via Cdk1.

## Introduction

The mitotic spindle must be precisely positioned in the cell not only to ensure accurate segregation of genetic material, but also to facilitate embryogenesis, cell fate determination, and tissue organization [1,2]. Defects in spindle orientation are thought to contribute to neurological diseases and cancer progression [3,4]. Depending on the cell type and environment, multiple intrinsic and extrinsic determinants (e.g. localization of polarity factors, or cell-cell contacts and other external mechanical cues) cooperate to determine the positioning of the spindle [5–7]. One of the first elements described to guide positioning of the spindle, during embryogenesis in amphibians, was cell shape [8]. Cell shape is also frequently observed to impact spindle orientation in cell lines plated as monolayers in tissue culture, where, as observed earlier in embryos, spindles are positioned during mitosis with respect to the long axis of the interphase cell [6,7,9]. In these single cell systems, it was later shown that extracellular adhesion forces from retraction fibers, which also contribute to cell shape, are transduced to the spindle to guide orientation [10,11]. It has recently become clear, however, that both cell shape and mechanical forces can separately provide cues for spindle orientation, and that spindle positioning in response to cell shape *per se* plays an important role in tissue and developmental contexts [12,13]. Because centrosomes are not always aligned with respect to the long cell shape axis (and retraction fiber forces) prior to mitosis, the spindle may need to “reorient”, or rotate by as much as 90 degrees in early mitosis in order to align with shape and adhesion cues [14].

Astral microtubules are essential for mitotic spindle orientation, including reorientation in response to cell shape [9,15,16]. They link the spindle to the cell cortex, where protein complexes interact with microtubules and provide the cues and forces required for positioning the spindle. The best characterized of these complexes contains the LGN, NuMA, and Gαi proteins, and cooperates with the microtubule motor dynein to link microtubules to the cell cortex and exert a pulling force [5,17]. The specific localization of such proteins can both dictate spindle positioning, for example in polarized systems or in response to adhesion patterns, as well as dynamically respond to the position of the spindle and chromosomes [18–20]. Given the central role of astral microtubules in spindle orientation, it is not surprising that many factors identified as important for spindle orientation promote the formation of astral microtubules, and when perturbed thus compromise the ability of astral microtubules to reach the cell cortex [21–29] [30]. While the precise regulation of microtubule dynamics has been shown to be critical for processes such as chromosome capture and alignment, or microtubule-kinetochore error correction [31], less is clear about the importance of astral microtubule dynamics to spindle positioning, or its precise regulation mechanisms. A few studies suggest that indeed more than the presence of astral microtubules at the cell cortex is required for spindle positioning. Depletion of Kinesin-13 or Kinesin-8 microtubule destabilizing kinesins, which results in longer, stabilized astral microtubules, has been shown to negatively impact spindle positioning [19,23,24,32–34]. Secondly, depolymerization of astral microtubules engaged with cortical NuMA/dynein complexes has been theorized to provide forces required for spindle positioning [1,35,36]. Pathways specifically regulating astral microtubule dynamics in time and space, however, remain unclear.

A dramatic increase in microtubule dynamics (i.e. shorter half-lives, increased catastrophes, and reduced rescues) accompanies the transition from interphase to mitosis, when the large interphase microtubule array is reorganized into a more compact mitotic structure [37–39]. This global increase in microtubule dynamics has been linked to activation of the Cdk1 kinase by Cyclin B, a major regulator of entry into mitosis [40–42], and is typically considered necessary for disassembling large interphase arrays, assembling the mitotic spindle, and aiding in search and capture of chromosomes. Less attention has been focused on how Cdk1-Cyclin B activity specifically modulates the properties of astral microtubules to facilitate spindle positioning [43].

Astral microtubule dynamics can be controlled by modulating microtubule nucleation and stability at the centrosome [26,44–46] and by impacting their dynamic plus-ends [21,47,48]. Major regulators of microtubule plus-end dynamics are the EB family of proteins, which are directly recruited to the growing plus-ends of microtubules [49]. EB1 is required for astral microtubule stability and spindle orientation, including shape-dependent positioning [1,21,22,50,51]. EB1 itself impacts microtubule dynamics, but can also directly recruit to the microtubule plus-ends a large number of microtubule plus-end tracking proteins (“+TIPS”), themselves necessary for several microtubule-dependent functions [49]. How distinct compositions of +TIPs are differentially recruited to EB proteins, to tune the dynamics and interactions of different microtubule populations for specific functions, including spindle orientation, is not well understood. Many +TIPs interact with EB1 via short conserved peptide motifs (SxIP) surrounded by basic, serine, and proline residues [52]. Phosphorylation of +TIPs close to these motifs appears to be a common mechanism to modulate their association with growing microtubule plus ends, by abolishing their interaction with EB1 [52–56]. However, Cdk1 phosphorylation has not been directly linked to its role in modulating microtubule dynamics at mitotic onset via a such a mechanism.

We previously found that a +TIP, GTSE1, is important for spindle orientation [23,53]. During mitosis GTSE1 inhibits the ability of the microtubule depolymerase MCAK to depolymerize microtubules, thus stabilizing them [23]. However, while GTSE1 interacts with microtubules and tracks microtubule plus-ends through interaction with EB1 in interphase, during mitosis it does neither [53]. Instead it is recruited to the mitotic spindle by direct interaction with spindle-associated clathrin heavy chain (CHC) [54]. GTSE1 is thus restricted to the inner spindle and spindle poles, where it controls astral microtubule length, likely via inhibition of centrosomal MCAK. This raises the question of why GTSE1 would be removed from all microtubules and EB1 at plus-ends in mitosis, only to be recruited back to inner spindles via a different mechanism. Here we show that the regulated loss of the microtubule stabilizing protein GTSE1 from microtubule plus ends at the onset of prometaphase is important to destabilize long astral microtubules to allow spindle orientation in response to cell shape. This is regulated by Cdk1, and required to allow destabilization of microtubules necessary for spindle reorientation in prometaphase. We have thus uncovered a direct mechanism by which Cdk1 promotes spindle orientation in early mitosis via astral microtubule destabilization.

## Results

### Cdk1-dependent phosphorylation of GTSE1 abolishes its interaction with EB1 and plus-end tracking during mitosis

GTSE1 associates with the microtubule lattice and growing microtubule plus ends in interphase, but this localization is reduced during mitosis [53]. To characterize the association of GTSE1 with growing microtubule plus-ends as cells enter mitosis, we imaged U2OS cells expressing GTSE1-GFP from a bacterial artificial chromosome (BAC) and a marker for microtubule plus-ends (EB3-mCherry), and quantified the relative intensities of both proteins at individual microtubule ends (Fig 1A, Supp Fig 1A). GTSE1 accumulated at EB3-positive microtubule plus ends in interphase, but was rapidly displaced from plus ends in mitosis, coincident with nuclear envelope breakdown (NEBD) (Fig 1A). In prometaphase and metaphase, GTSE1 localized to the inner-spindle, which we previously reported is via interaction with spindle-associated clathrin [54]. Upon anaphase onset, GTSE1 resumed its localization at growing microtubule plus ends (Supp Fig 1A). To establish whether the loss of GTSE1 plus-end tracking during mitosis was due to loss of interaction with EB1, we used purified EB1-GST to pull down GTSE1 from U2OS cell extracts arrested in either S-phase or prometaphase. GTSE1 from S-phase, but not prometaphase, extract interacted with EB1-GST, confirming the loss of this interaction in mitosis (Supp Fig 1B).

**Figure 1:**
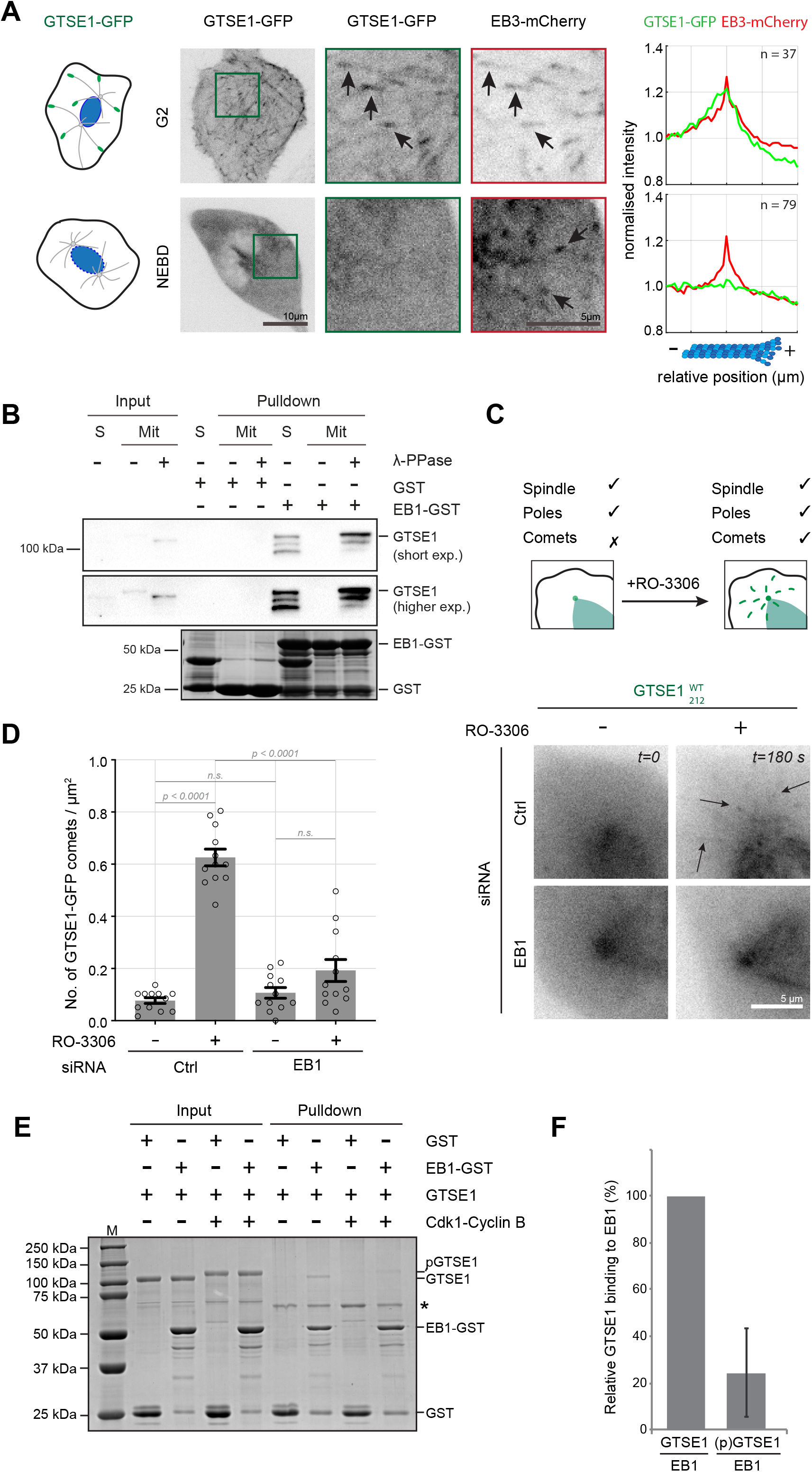
Cdk1 phosphorylates GTSE1 and prevents its interaction with EB1 and microtubule plus-end tracking in mitosis. **(A)** Schematic depicting GTSE1 localization at microtubule plus ends in G2 and at nuclear envelope breakdown (NEBD). Live cell images of U2OS cells stably expressing GTSE1-GFP from a BAC (green) and EB3-mCherry from a plasmid (red). Scale bar 10 μm. Magnified insets show GTSE1 (middle) and EB3 (right) with arrows indicating comets, scale bar 5 μm. Averaged line scan intensity profiles of microtubule plus-ends show relative intensities of GTSE1-GFP (green) and EB3-mCherry (red), centered at the maximum EB3 intensity. Line scan insets indicate the number of microtubule plus ends from 2 cells in 2 experiments expressing comparable levels of both transgenes. **(B)** Disruption of the EB1-GTSE1 interaction during mitosis requires phosphorylation. Immunoblot (anti-GTSE1) of an EB1-GST pulldown using U2OS cells arrested in S-phase or prometaphase, the latter also treated (or not) with λPPase. Below is Coomassie Blue staining of GST and EB1-GST pulled down in all three conditions. **(C)** Inhibition of Cdk1 restores EB1-dependent GTSE1 plus-end tracking in mitosis. Schematic depicting GTSE1 localization (in green) on the spindle, poles and comets in early prometaphase before and after addition of Cdk1 inhibitor (RO-3306). Stills from live cell microscopy of U2OS cells stably expressing GTSE1-GFP (U2OS^WT^_212_) before and after addition of RO-3306. U2OS^WT^_212_ cells were transfected with Control (Ctrl) or EB1 siRNA for 48 hours. Cells were imaged at 1 second-intervals for 20 seconds in prometaphase and then treated with RO-3306 and imaged similarly for 3 minutes. Arrows indicate comets. See Videos 1 and 2. Scale bar 5 μm. **(D)** Quantification of number of GFP comets per unit area in the four conditions shown in C. n = 12 cells per condition over 2 experiments. P-values from unpaired two-tailed t-test using Welch’s correction. Error bars indicate s.e.m. **(E)** Cdk1 phosphorylation of GTSE1 reduces interaction with EB1 *in vitro*. Coomassie Blue stained gel of a GST pulldown assay using EB1-GST as bait to pull down purified GTSE1 phosphorylated (or not) with Cdk1-Cyclin B., ‘*’ indicates GSH beads blocked using BSA. **(F)** Quantification of relative GTSE1 binding to EB1 before and after phosphorylation of GTSE1 by Cdk1. n = 3 experiments. Error bars represent s.d.

We next sought the mechanism by which cells control the dissociation of GTSE1 from EB1 in mitosis. We first confirmed that GTSE1 is hyperphosphorylated during mitosis in human cells, as has been shown in mouse and *Xenopus* (Supp Fig 1C) [53,55]. We next asked whether this affected the GTSE1-EB1 interaction by performing an EB1-GST pulldown as above using mitotic lysates, but after treatment with λ-phosphatase to dephosphorylate mitotic proteins. GTSE1 from dephosphorylated mitotic lysate could be pulled down by EB1, suggesting that this interaction is regulated by mitotic-specific phosphorylation (Fig 1B). To identify the kinase(s) responsible for disrupting the GTSE1 interaction with EB1 in mitosis, we asked whether inhibition of different mitotic kinases with small molecular inhibitors could force GTSE1 to localize to growing microtubule plus ends in mitosis. U2OS cells expressing GTSE1-GFP from a BAC transgene (GTSE1^WT^_212_; [53]) were thus imaged in mitosis after treatment with inhibitors against Aurora A (MLN8054), Aurora B (ZM447439), Plk1 (BI 2536), or Cdk1 (RO-3306) [56–59]. No discernable plus-end tracking of GTSE1-GFP was observed in mitosis in untreated cells (Fig 1C, Supp Fig 1D), nor after inhibition of Aurora A, Aurora B, or Plk1 (Supp Fig 1D). Inhibition of Aurora A did cause complete displacement of GTSE1 from the mitotic spindle, as previously reported [60]. In contrast, inhibition of Cdk1 shortly after NEBD led to the appearance of moving comet-like structures of GTSE1-GFP, similar to that seen in interphase (Fig 1C; Video 1). Quantification of the number of detectable GTSE1-GFP comets in mitosis indicated an approximately 6-fold increase after Cdk1 inhibition (Fig 1D). Washing out the Cdk1 inhibitor 30 seconds later caused these comets to rapidly disappear again (Video 2). Quantification of GTSE1-GFP levels at EB3-mCherry-labelled plus-ends in a dual-labeled cell line indicated that 90% of these plus-ends now accumulated GTSE1 after Cdk1 inhibition (Supp Fig 1E, F). All Cdk1-inhibition experiments were performed within a few minutes of NEBD to avoid cells exiting mitosis during analysis. To confirm that the relocalization of GTSE1 to growing microtubule plus-ends was dependent on EB1, we repeated these experiments after EB1-depletion (Fig 1C and D, Supp Fig 1G; Video 3). Following RNAi of EB1, we no longer observed an increase in the number of GTSE1-GFP-positive comets after Cdk1 inhibition. To test whether phosphorylation of GTSE1 by Cdk1 directly disrupts its interaction with EB1 *in vitro*, we performed pulldown experiments with purified proteins after phosphorylation by Cdk1-CyclinB. EB1-GST was incubated with either unphosphorylated or Cdk1-phosphorylated GTSE1. Phosphorylation of GTSE1 reduced its interaction with EB1 by 80% (Fig 1E and F). In contrast, phosphorylation of EB1 by Cdk1 did not affect its interaction with GST-GTSE1, demonstrating that this interaction is regulated by the action of Cdk1 on GTSE1 (Supp Fig 1H and I). Thus, Cdk1 removes GTSE1 from growing microtubule plus-ends during mitosis by disrupting its interaction with EB1, constraining GTSE1 to the clathrin-associated inner-spindle.

### Mutating Cdk1- and mitosis-dependent phosphorylation sites around its EB1-interaction motifs causes GTSE1 to plus-end track in mitosis

We next aimed to identify the phosphorylation sites on GTSE1 that regulate the interaction with EB1. First, we identified mitosis-specific phosphorylation sites on GTSE1 by immunoprecipitating it from U2OS cells arrested in either S-phase or prometaphase, and subjecting it to mass spectrometry analysis. This revealed a total of 29 mitosis-specific phosphosites on GTSE1 from a total peptide coverage of 70% (Fig 2A, Table I). 27 of these 29 sites were previously identified in large scale proteomic studies in mitotic cells (Fig 2A and Table I) [61–63]. Many of the mitosis-specific phosphorylation sites identified by us and others were concentrated in a region 20 residues upstream and downstream of the two SxIP motifs, suggesting that phosphorylation at these sites could potentially hinder electrostatic interactions between GTSE1 and EB1 (Fig 2A). We next purified His-tagged GTSE1, phosphorylated it *in vitro* with recombinant Cdk1-Cyclin B (Supp Fig 2A), and analyzed phosphorylated residues by mass spectrometry. 55 sites on GTSE1 were phosphorylated by Cdk1 *in vitro*. Combining all mitotic and Cdk1-dependent phosphorylation data, we identified 14 phosphorylation sites within a 58-amino acid stretch surrounding the two SxIP sites on GTSE1 (Fig 2A).

**Figure 2:**
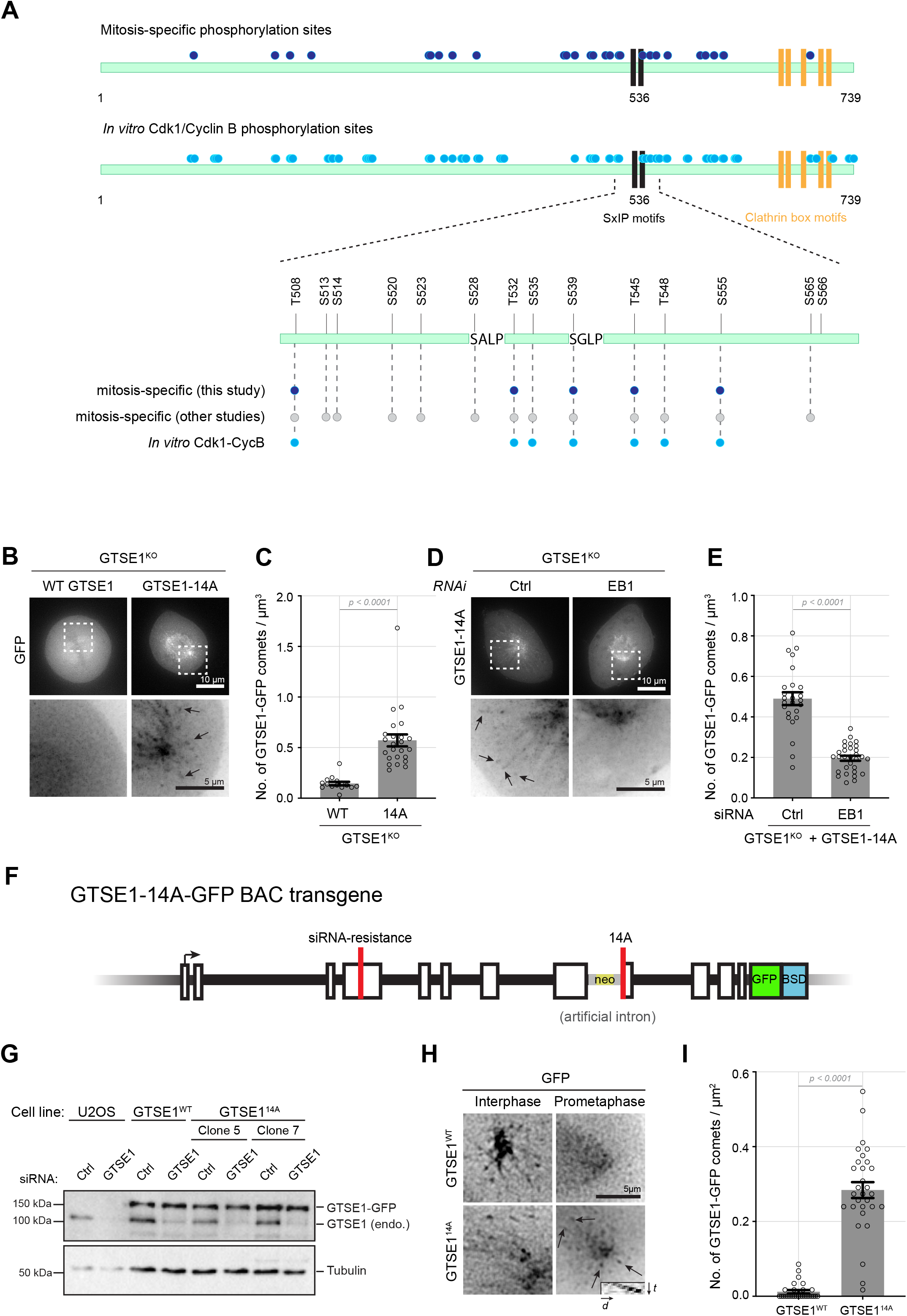
A phospho-mutant of GTSE1 plus-end tracks in mitosis in an EB1-dependent manner. **(A)** Mitotic and Cdk1 phosphorylation sites identified in this study in GTSE1. Schematic representation of GTSE1, with two SxIP motifs shown in black and clathrin box motifs in yellow. Mitosis-specific (dark blue circles) and *in vitro* Cdk1-Cyclin B (light blue circles) phosphosites are shown. The 58 amino acid region between 508-566 on GTSE1 is shown in more detail, with mitosis-specific phosphorylation sites identified by others in high throughput MS studies (gray circles) additionally represented. **(B)** Single frames from live imaging in mitosis of U2OS cells knocked out for GTSE1 (GTSE1^KO^) cells transiently transfected with plasmids expressing wildtype GTSE1-GFP or GTSE1-14A-GFP. Scale bar 10 μm. Magnified insets show comets indicated by arrows in cell expressing GTSE1-14A. Scale bar 5 μm. See Video 4. **(C)** Quantification of number of GFP comets per unit volume from experiment shown in (B). n = 16-24 cells from 3 independent experiments. P-value from unpaired t-test with Welch’s correction. **(D)** Single frames from live imaging in mitosis of GTSE1^KO^ cells treated with Control (Ctrl) or EB1 siRNA for 48 hours followed by transient transfection of plasmids expressing GTSE1-14A-GFP. Scale bar 10 μm. Magnified insets show comets indicated by arrows. Scale bar 5 μm. See Video 5. **(E)** Quantification of number of GTSE1-14A-GFP comets per unit volume in presence and absence of EB1 from experiment shown in (D). n = 25-30 cells from 3 independent experiments. P-value from unpaired t-test with Welch’s correction. **(F)** A schematic showing the design of the GTSE1-14A-GFP BAC containing exons and introns and mutations for resistance to siRNA. The intron between exons 8 and 9 and the wildtype exon 9 was replaced with an artificial intron that confers neomycin resistance and the mutated exon 9 with 14 Ser/Thr residues mutated to alanine. **(G)** Western blot showing comparable levels of BAC-expressed GFP tagged wildtype (GTSE1^WT^) or two independent clones expressing GTSE1-14A transgene (GTSE1^14A^), before and after depletion of endogenous GTSE1 for 48 hours using siRNA. Immunoblot using anti-GTSE1 and anti-tubulin antibodies. **(H)** Stills from live microscopy of U2OS cells stably expressing wildtype GTSE1 (GTSE1^WT^) or GTSE1-14A (GTSE1^14A^) in interphase and prometaphase. Arrows indicate comets in mitosis in GTSE1^14A^ and the inset shows a kymograph of a GTSE1-14A labeled microtubule plus end. Scale bar 10 μm. See Video 6. **(I)** Quantification of mean number of GFP comets per unit area in prometaphase from experiment shown in (H). At least 28 cells of GTSE1^WT^ and GTSE1^14A^ were analyzed from 3 independent experiments. P-value from unpaired t-test with Welch’s correction. Error bars indicate s.e.m. unless otherwise indicated.

To evaluate which phosphorylation sites in the proximity of the EB1-interaction motifs in GTSE1 disrupted EB1-interaction during mitosis, we generated a series of GTSE1-GFP mutants with these phosphorylation sites mutated to alanine (Supp Fig 2B). The mutants were then transiently transfected into GTSE1 knockout cells (GTSE1^KO^; [23]) and their ability to plus end-track in mitotic cells was evaluated by live microscopy. We initially mutated two conserved Cdk1 consensus sites located directly adjacent to the SxIP motifs (GTSE1^TP/AA^). This mutant however displayed no apparent plus-end tracking in mitosis (Supp Fig 2B). We then constructed several mutants with increasing numbers of mutated phosphorylation sites. While all mutants plus-end tracked in interphase, we could only observe weak tip-tracking in mitosis after mutating 7 phosphosites, and found that mutation of all 14 phosphosites resulted in the most robust plus-end tracking (Fig 2B and C, Supp Fig 2B; Video 4). To confirm this mutant was plus-end tracking via EB1 interaction, we depleted EB1 in the presence of the mutant, which indeed abolished the increased plus-end tracking (Fig 2D and E, Supp Fig 2C; Video 5). Thus, disrupting mitotic phosphorylation of GTSE1 prevents removal of GTSE1 from microtubule plus-ends at mitosis.

To study the importance of the disruption of the EB1-GTSE1 interaction for mitotic function, we generated the 14 phosphosite mutations in an RNAi-resistant BAC harbouring GTSE1-GFP in a single step via ESI mutagenesis (Fig 2F; [64]). We then generated stable and clonal cell lines expressing RNAi-resistant versions of either wildtype GTSE1-GFP or the GTSE1-14A-GFP mutant (GTSE1^WT^ and GTSE1^14A^) (Fig 2G). Both transgenes exhibited plus-end tracking in interphase (Video 6). Additionally, GTSE1-14A-GFP expressed at near endogenous levels was clearly recruited to growing microtubule plus-ends in mitosis, unlike wildtype GTSE1-GFP (Fig 2H and I; Video 6).

### Restoring GTSE1 plus-end tracking during mitosis increases astral microtubule length

In interphase, GTSE1 tracks microtubule plus-ends and stabilizes microtubule growth duration [53]. We thus analyzed whether microtubule stability in mitotic GTSE1^14A^ cells was changed as compared to wildtype cells. Fixed imaging of mitotic GTSE1^14A^ cells revealed an increased abundance of astral microtubules, that appeared to often reach the cell cortex (Fig 3A). We then used EB1 comet signals and their distances from the centrosome to quantify the number and individual lengths of astral microtubules [47,54]. As previously shown, depletion of GTSE1 reduces both the number and length of metaphase astral microtubules, and an RNAi-resistant GTSE1-GFP transgene (GTSE1^WT^) rescues these defects (Fig 3A-C) [23,54]. Cells expressing equivalent protein levels of GTSE1-14A, however, displayed significantly longer and more abundant astral microtubules in metaphase, even before RNAi-depletion of endogenous GTSE1 (Fig 3A-C). We verified this effect with an independently isolated cell line generated with the same mutant BAC (Supp Fig 3A and B). This indicated a dominant effect of the mutant, which was expected given the gain of function of EB1-dependent microtubule plus-end localization. This stabilization effect of the mutant appeared specific to astral microtubules, as there was not a significant change in the number of EB1 comets within the spindle as compared to U2OS or GTSE1^WT^ cells (Fig 3D), and the total tubulin intensity within the inner spindle was not affected (Supp Fig 3C and D). Plotting the distribution of metaphase astral microtubule lengths among cell lines suggested that in the GTSE1^14A^ mutant, there was a selective increase in the number of longer (6-10 μm) astral microtubules (Fig 3C). Indeed, plotting the fold difference in astral microtubule numbers at different lengths showed no significant difference of microtubules less than 6 μm long, nearly 2-fold as many microtubules 6-9 μm long, and over 3-fold more microtubules longer than 9 μm in GTSE1^14A^ cells (Supp Fig 3E).

**Figure 3:**
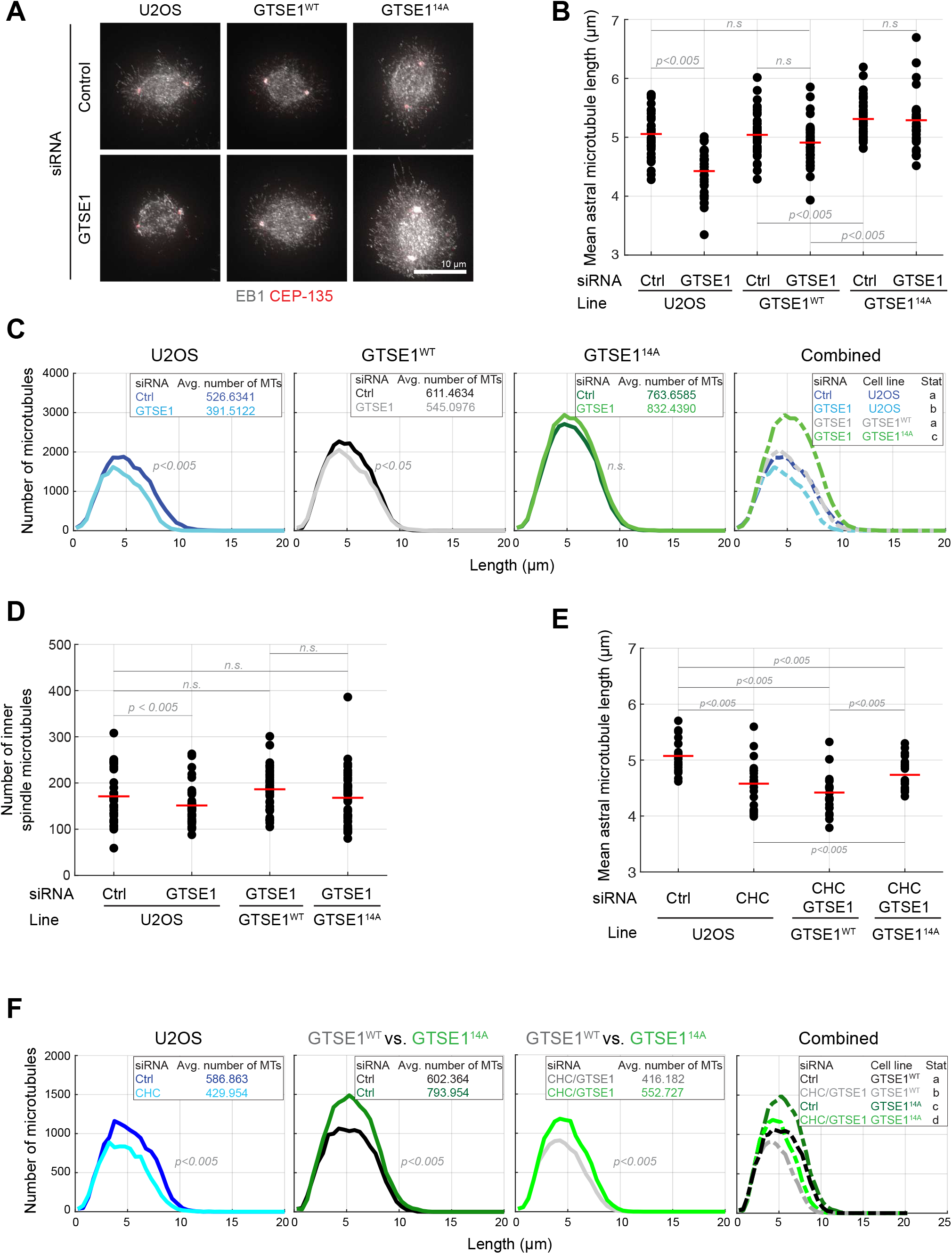
Restoring GTSE1 to growing microtubule plus-ends in mitosis stabilizes astral microtubules in a clathrin-independent manner. **(A)** Immunofluorescence images in metaphase of U2OS, GTSE1^WT^ and GTSE1^14A^ cells treated with control (Ctrl) or GTSE1 siRNA. Cells were fixed and stained with antibodies against EB1 (in gray) and CEP-135 (in red). Scale bar 10 μm. **(B)** Quantification of mean astral microtubule length from experiment shown in (A). The red horizontal line indicates mean. n = 37 cells from 3 independent experiments, pooled for representation. P-values from unpaired two-tailed student’s t-test. **(C)** Distributions of astral microtubule numbers as a function of their lengths. n = 37 cells from 3 independent experiments, pooled for representation. The mean number of astral microtubules is presented in insets. P-values from unpaired two-tailed student’s t-test. Inset in ‘Combined’ shows the significances (different letters indicate groups differing significantly). **(D)** Quantification of number of EB1 labeled microtubules in the inner spindle in metaphase U2OS, GTSE1^WT^ and GTSE1^14A^ cells after control (Ctrl) or GTSE1 RNAi. The mean is shown in red. P-values from unpaired two-tailed student’s t-test. **(E)** Quantification of mean astral microtubule lengths from 3D reconstructions of cells images in metaphase of U2OS, GTSE1^WT^ and GTSE1^14A^ cells treated with control (Ctrl), CHC siRNA alone (U2OS) or a combination of CHC and GTSE1 siRNA (in GTSE1^WT^ and GTSE1^14A^). The mean is marked in red. n = 22-43 cells from 3 independent experiments, pooled for representation. P-values from unpaired two-tailed student’s t-test. **(F)** Distributions of astral microtubule numbers as a function of their lengths. n = 22 cells from 3 independent experiments, pooled for representation. The mean number of astral microtubules is presented in insets. P-values from unpaired two-tailed student’s t-test. Inset in ‘Combined’ shows the significances (different letters indicate groups differing significantly).

GTSE1-mediated stabilization of astral microtubules in wildtype cells is clathrin-dependent, requiring clathrin-mediated recruitment of GTSE1 to the spindle poles and inner-spindle [54]. Due to the localization of GTSE1-14A protein to astral microtubule plus-ends and the dominant impact on astral length, we hypothesized the gain-of-function of this mutant would be clathrin-independent. To test this, we depleted CHC in GTSE1^WT^ and GTSE1^14A^ cells (Supp Fig 3F). Indeed, even after removal of CHC, the GTSE1-14A mutations remained able to induce longer astral microtubules, supporting a clathrin-independent gain of GTSE1 function at astral microtubule plus-ends (Fig 3E and F, Supp Fig 3G). Furthermore, CHC still contributed to astral microtubule length in GTSE1^14A^ cells (Fig 3F, compare dark green and light green traces), again indicating separate pathways.

As Cdk1-mediated removal of GTSE1 from growing microtubule plus-ends occurs at NEBD, we next asked whether the defect in the ability of the mitotic GTSE1^14A^ cells to control astral microtubule length also manifested in prometaphase. Quantification of microtubule lengths and abundance from asters in early prometaphase cells revealed a significantly increased number of astral microtubules, which was again characterized by an apparent increase in the number of longer astral microtubules (Fig 4A-C). Plotting the fold difference in astral microtubule numbers of different lengths in prometaphase cells showed no significant difference of microtubules less than 8.5 μm long, yet over 3-fold more microtubules longer than 8.5 μm in GTSE1^14A^ cells compared to GTSE1^WT^ (Fig 4D). For insight into the underlying changes leading to longer astral microtubules, we analyzed microtubule dynamics in prometaphase cells additionally expressing mCherry-EB3. While we did not observe any change in microtubule growth rates in GTSE1^14A^ cells compared to GTSE1^WT^, the average catastrophe frequency reduced (Fig 4E and F). Thus, the GTSE1-EB1 interaction is negatively regulated by phosphorylation in early mitosis to confine the number of long astral microtubules, likely by increasing the chance of microtubule catastrophe.

**Figure 4:**
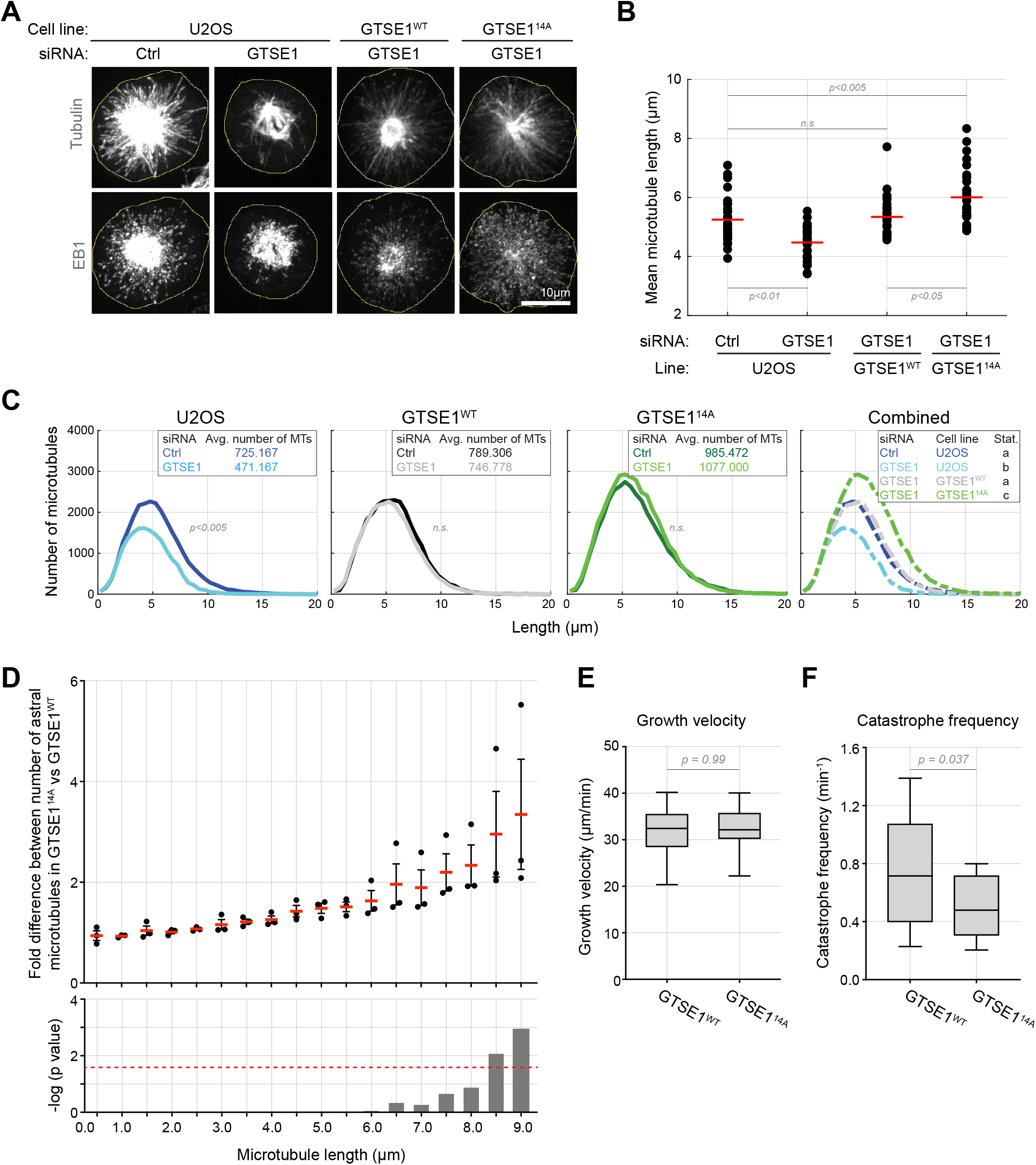
CDK1-mediated removal of GTSE1 from microtubule plus ends destabilizes long astral microtubules and promotes microtubule catastrophe in prometaphase. **(A)** Immunofluorescence images in prometaphase of U2OS, GTSE1^WT^ and GTSE1^14A^ cells treated with control (Ctrl) or GTSE1 siRNA. Cells were fixed and stained with anti-tubulin and anti-EB1 antibodies. Scale bar 10 μm. **(B)** Quantification of mean microtubule lengths from experiment shown in (A). Red line indicates mean. n = 43 cells over 3 independent experiments. P-values from unpaired two-tailed student’s t-test. **(C)** Distributions of microtubule numbers as a function of their lengths. n = 43 cells from 3 independent experiments, pooled for representation. The mean number of microtubules is presented in insets. P-values from unpaired two-tailed student’s t-test. Inset in ‘Combined’ shows the significances (different letters indicate groups differing significantly). **(D)** Quantification of the fold difference in the number of microtubules of different lengths in prometaphase between GTSE1^WT^ and GTSE1^14A^. P-values from one-way ANOVA. The horizontal red dashed line indicates a p-value of 0.05. **(E)** Mean microtubule growth velocity in mitosis in GTSE1^WT^ and GTSE1^14A^ cells, obtained from movies of cells co-expressing EB3-mCherry, analyzed with plusTipTracker software. n = 17-19 cells from one experiment. P-values from unpaired t-test with Welch’s correction. **(F)** Mean catastrophe frequency in GTSE1^WT^ and GTSE1^14A^ cells co-expressing EB3-mCherry calculated using plusTipTracker software. n = 10-14 cells from one experiment, only values lying between 10^th^ and 90^th^ percentiles were considered. P-values from unpaired t-test with Welch’s correction. Error bars represent s.e.m. unless otherwise indicated.

### Phospho-dependent displacement of GTSE1 from microtubule plus ends in prometaphase is required for spindle reorientation via astral microtubule destabilization

Given the importance of astral microtubules for spindle reorientation, we asked whether restoring the mitotic EB1-GTSE1 interaction and subsequent hyperstabilization of astral microtubules led to a defect in spindle orientation that could explain the need to disrupt this interaction. We previously reported that cells depleted of GTSE1 are defective in metaphase spindle orientation with respect to the substrate due to shorter astral microtubules. We thus asked whether the longer metaphase astral microtubules observed in GTSE1^14A^ cells also perturbed this type of orientation. While the defect after GTSE1 depletion was restored in GTSE1^WT^ cells, GTSE1^14A^ cells were defective, suggesting the failure of these cells to destabilize long astral microtubules compromised spindle orientation in metaphase (Supp Fig 4A). Loss of GTSE1 causes an increased mitotic duration due to chromosome congression defects [23,54,65]. As chromosome congression defects can perturb spindle orientation [20], we analyzed mitotic timing in GTSE1^14A^ cells. We did not observe any defect, suggesting they are proficient for chromosome alignment and that this could not explain the spindle orientation defects (Supp Fig 4B).

We next assayed the ability of the prometaphase spindles in GTSE1^14A^ cells to orient their axis relative to cell shape/adhesion cues by imaging cells with microtubules labeled entering and proceeding through mitosis. The initial cell shape axis was determined 8 minutes prior to NEBD, and compared to the centrosome-to-centrosome spindle axis at anaphase onset. When considering cells in which the centrosome-centrosome axis at NEBD was more than 30 degrees deviated from the cell shape axis (and thus would require active repositioning), approximately 53% of U2OS cells were able to reorient spindles to match the shape axis (Fig 5A and B). Depletion of GTSE1, which destabilizes astral microtubules, severely compromised this reorientation (13% reoriented), while the presence of a RNAi-resistant wildtype GTSE1 restored it (58%). Strikingly, GTSE1^14A^ cells were also severely compromised for spindle reorientation (20%), suggesting that stabilization of astral microtubules can perturb the ability of cells to reorient spindles relative to cell shape. Although the defect in the final orientation of cells both lacking, and with longer astral microtubules (GTSE1 RNAi in U2OS, and GTSE1^14A^, respectively) were comparable, we noticed that spindles in the former tended to randomly rotate within the cell, while the latter appeared more static than normal. To quantify these observations, we determined the difference between the final orientation of the spindle (α_Anaphase_) and the initial orientation of the spindle (α_NEBD_), relative to the interphase shape axis (again in cells with an initial α_NEBD_ of more than 30 degrees; Fig 5C). As expected, spindles in both control cells and GTSE1^WT^ cells corrected their orientation, indicated by the angle becoming smaller (Fig 5C). In cells depleted of GTSE1, spindle orientation angles actually became more deviant than they started, consistent with a lack of microtubules resulting in a complete failure to sense cell shape. In contrast, spindles in GTSE1^14A^ cells on average did not change from their initial, off-axis position, suggesting that they resisted rotation and were “stuck” at their pre-prometaphase position. As this defect could arise from excessive and/or stabilized engagement of cortical astral microtubule-interacting complexes, we measured the number of astral microtubules reaching the cell cortex in prometaphase (Fig 5D). Consistently, cells expressing GTSE1-14A contained more than 3 times the number of astral microtubules reaching the cortex than under wildtype conditions. Together, these results indicate that mitotic phosphorylation of GTSE1, which disrupts the interaction with EB1 and microtubule plus-end tracking, is required for spindle orientation. Cells containing mutations disrupting this regulation are unable to reorient the mitotic spindle in response to cell shape, which we attribute to the coincident stabilization of long astral microtubules reaching the cell cortex.

**Figure 5:**
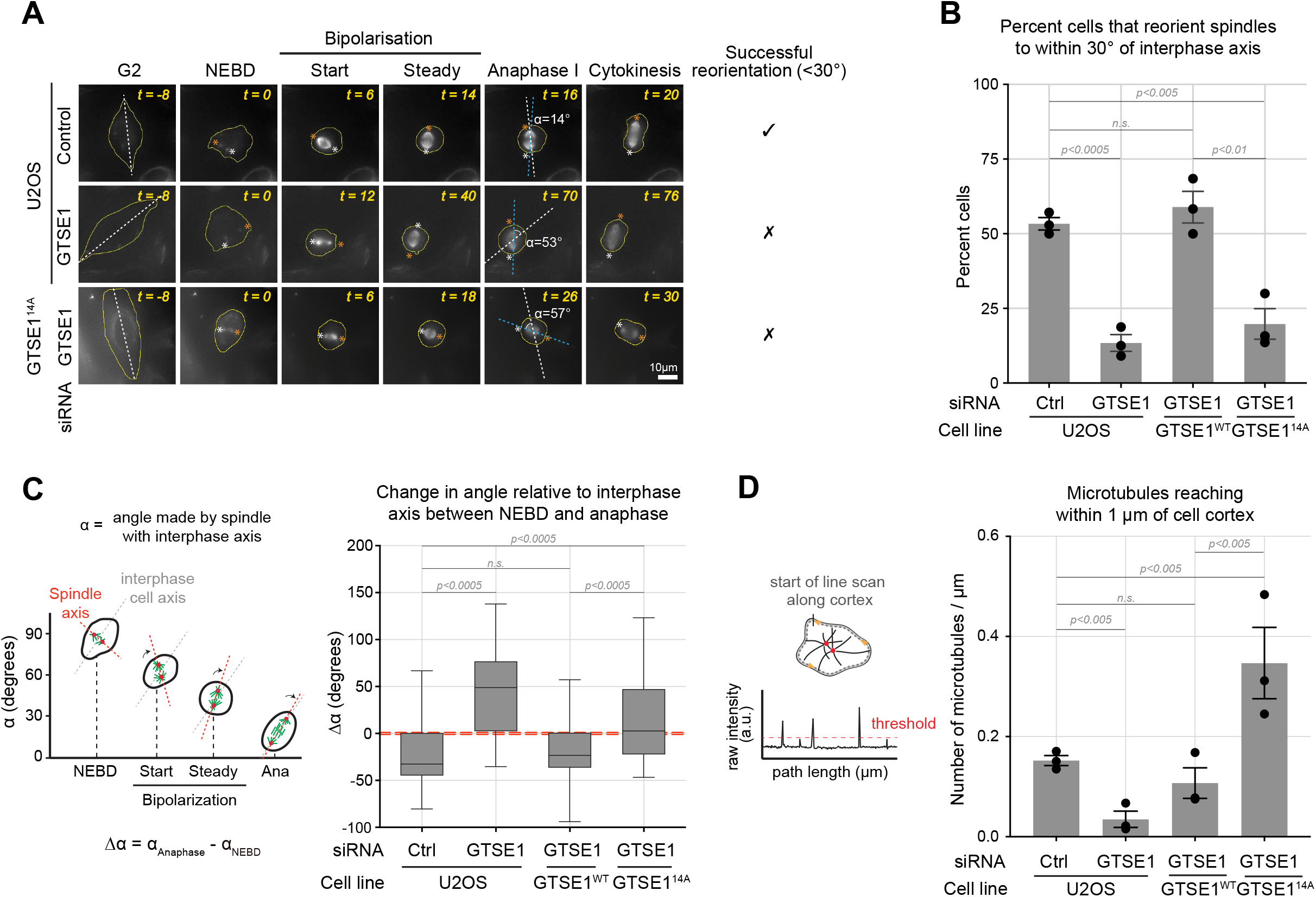
Restoring GTSE1 to growing microtubule plus-ends in mitosis inhibits the ability of cells to reorient spindles in response to cell shape cues. **(A)** Stills from live-cell imaging of U2OS and GTSE1^14A^ cells treated with control (Ctrl) or GTSE1 siRNA for 48 hours and imaged in presence of the cell-permeable siR-Tubulin dye. Individual centrosomes are labeled with ‘*’ in white and orange. Dashed white lines indicate the cell shape axis, dashed blue lines indicate the spindle axis. Scale bar 10 μm. **(B)** Quantification of spindle positioning defect in experiment shown in (A). Only cells that have their spindle axis more than 30° away from the interphase axis at NEBD are considered for analysis. Percent cells that are able to align their spindles to within 30° of the interphase axis is shown (with individual experimental means displayed). n = 39-51 cells over 3 experiments. P-values from unpaired t-test. **(C)** Left panel: Schematic illustrating the change in angle α made by the spindle with the interphase axis at different phases as indicated on the graph. An example illustrating progressive spindle rotation to eventually align along the interphase axis is shown. A negative value for Δα indicates spindle movement towards long axis, while a positive value indicates spindle movement away from the long axis. Right panel: Box plot of Δα values in U2OS, GTSE1^WT^ and GTSE1^14A^ cells. P-values from unpaired t-test with Welch’s correction. **(D)** Left panel: Schematic illustrating the method for semi-automatic detection of microtubules reaching the cortex. Quantification of the number of microtubules reaching within 1 μm of the cell cortex, per μm of cortex in 2-dimensions in U2OS, GTSE1^WT^ and GTSE1^14A^ prometaphase cells (with individual experimental means displayed). n = 30-37 cells over 3 independent experiments. P-values from unpaired t-test. All error bars represent s.e.m. unless otherwise indicated.

## Discussion

Despite the importance of astral microtubules to mitotic spindle orientation, little is known about how their dynamics and stability may be specifically controlled to contribute to this process. Here we have shown that destabilization of long astral microtubules in prometaphase is required for spindle orientation in response to cell shape. This destabilization is controlled at NEBD by Cdk1 phosphorylation of GTSE1, which disrupts the recruitment of GTSE1 to growing microtubule plus-ends by abolishing its interaction with EB1. Loss of GTSE1 from microtubule plus-ends in mitosis promotes their instability by raising their catastrophe frequency.

We have thus described a new mechanism by which Cdk1 controls spindle orientation in response to cell shape. While Cdk1 activity has been previously linked to spindle orientation via negatively regulating the cortical localization of NuMA [66], we show the first link via regulation of microtubule stability. Cdk1 activity has long been known to promote increased microtubule dynamics as cells transition to mitosis [40–42]; to our knowledge this is the first example of a mechanism linking Cdk1 phosphorylation of a target protein to specifically control astral microtubule stability in metazoans. Targeting an EB1-+TIP interaction may be a repeated mechanism for the Cdk1-dependent regulation of microtubule dynamics that happens in mitosis. Although only a few +TIPs have been reported to be removed from microtubules in mitosis [67–69], we have identified via quantitative mass spectrometry more than 10 SxIP-containing proteins that interact with EB1 in interphase, but not mitosis (unpublished data). Some have likely been previously overlooked due to commonly used overexpressing GFP-transgene constructs. Indeed, when dramatically overexpressed, GTSE1-GFP plus-end tracks during mitosis (unpublished observation), probably due in part to overwhelming the cellular phosphorylation machinery.

While loss of EB1 (and thus the associated array of +TIPs from growing ends) has been documented to inhibit microtubule dynamics and destabilize astral microtubules [21,22,70], our work here highlights that there is a dynamic and precise tuning of astral microtubule length and dynamics mediated by the spatiotemporal regulation of +TIP interactions. Perturbations causing dramatic global hyperstabilization microtubules throughout the cell, such as treatment with Taxol, are known to affect spindle positioning [71,72]. What we show here is that even relatively subtle changes to astral microtubule stability nevertheless have a quite dramatic impact on spindle repositioning in prometaphase and thus must be precisely controlled. Our results suggest that the spindle orientation defect exhibited by GTSE1^14A^ cells arises from hyperstabilization of long astral microtubules in prometaphase. This is consistent with the observation that perturbation of the microtubule destabilizing kinesin Kif18B, which also results in longer astral microtubules, leads to spindle orientation defects [19,33,47,73,74]. Interestingly, while GTSE1 is removed from microtubule plus ends during mitosis, Kif18B is specifically recruited in mitosis to astral microtubule plus ends by EB1 [47,73], highlighting the importance of regulating astral microtubule length and dynamics via modulation of +TIP-EB1 interactions.

Why must long astral microtubules be destabilized in prometaphase? We show that in stark contrast to cells lacking astral microtubules, which display exaggerated and random spindle movements, cells with hyperstabilized long astral microtubules resist spindle rotation in response to cell shape. Under these conditions, a combination of excessive engagement of cortical microtubule-interaction sites and a reduced likelihood of a catastrophe/depolymerization event could pose a significant barrier to reorientation of a spindle. Excessive longer astral microtubules may establish too many cortical interactions early in prometaphase, and in combination with a lack of microtubule dynamicity, inhibit the ability of spindles to dynamically react to cell shape. This is consistent with a simple model from Minc et al (2011) of how microtubules can position nuclei by sensing cell geometry, which was also applied to spindle reorientation in response to cell shape [15]. In this model, microtubules “probe” the cell shape and exert length-dependent pulling when engaged with force generators at the cell cortex. Consistently, increasing the numbers of microtubules reaching the cortex in the first division of *C. elegans* embryos increases the “centering stiffness” of the spindle, or force required to displace the spindle within the cell [32]. Additionally, evidence suggests that microtubules may need to be dynamic at the cell cortex in order to generate depolymerization-coupled force generation, for example via NuMa, to position spindles [35,36,72,75].

We show that Cdk1 destabilizes long astral microtubules by removing the microtubule-stabilizing protein GTSE1 from growing microtubule plus-ends. GTSE1 stabilizes microtubules by inhibiting the microtubule depolymerization activity of MCAK, which also localizes at astral microtubule plus-ends during mitosis [23,78]. The decreased catastrophe frequency and longer lengths of microtubules observed when GTSE1 is not removed from astral microtubule plus-ends (i.e. in GTSE1^14A^ cells) thus likely reflects inhibition of the microtubule plus-end associated pool of MCAK. Indeed, the reduction in catastrophe frequency we observe in GTSE1^14A^ mutant cells is similar to that observed when MCAK is in inhibited in PtK2 cells [76]. Consistently, inhibition of MCAK by RNAi leads to longer astral microtubules and spindle positioning defects [23,24]. In the future, molecular definition of the MCAK inhibition mechanism of GTSE1 will allow point mutation of GTSE1-14A to test this directly. GTSE1 normally stabilizes astral microtubules when recruited to spindle poles by clathrin, likely through inhibition of centrosomal MCAK [54]. However, we have shown here that the ability of GTSE1-14A mutant to stabilize astral microtubules is independent of clathrin, consistent with it inhibiting a distinct pool of MCAK at growing microtubule plus-ends.

In summary, we have provided evidence here that astral microtubule dynamics must be precisely regulated to ensure spindle orientation mechanisms, in particular that long astral microtubules must be selectively destabilized. We further have elucidated a molecular pathway by which these changes are both spatially and temporally regulated by Cdk1 activity at the onset of prometaphase: by disrupting recruitment of the microtubule stabilizing protein GTSE1 to microtubule plus ends via EB1. Regulation of EB1-+TIP complexes may be a common theme by which Cdk1 induces the substantial changes in microtubule dynamics accompanying mitosis.

## Acknowlegdements

We thank A. Musacchio for help with reagents, and P. Singh and C. Körner for purified Cdk1-CyclinB. We thank M. Tapaswi for help with MATLAB analyses. We thank G. Vader, A. Rondelet, and H. Thakur for comments on the manuscript. The authors declare no competing financial interests. This work was supported by the Max Planck Institute of Molecular Physiology.

## Materials and Methods

### Cloning and plasmids

Plasmids for mammalian expression were derived from the pEGFP-N2 GTSE1-GFP-T2A-Bsd, a previously modified version [23] of the pEGFP-N2 vector (Addgene). Mutations were introduced into the GTSE1 cDNA by either site-directed mutagenesis or Gibson cloning, with the exception of the GTSE1^TP/AA^ BAC that was previously generated in the laboratory. The GST-hGTSE1 construct used for protein purification from insect cells was also described in the study mentioned above. The GTSE1-14A BAC carrying point mutations at T508, S513, S514, S520, S523, S523, S528, T532, S535, S539, T545, T548, T555, S565, S566 was generated through recombineering by ESI mutagenesis based on Redβ and Redγ expression [64]. Briefly, a cassette was first created that contains mutated exon 9 linked via a T2A peptide to neomycin resistance gene flanked by splice sites at both ends. This cassette was amplified by PCR using primers that provide 50 bp long homology arms to specify the site of recombination on the GTSE1 WT BAC. Standard recombination based on inducible Redβ and Redγ expression was carried out and positive clones were tested for the correct recombination event. The integrity of GTSE1-14A BAC was checked by performing restriction digestion with NotI (New England Biolabs) and comparing the obtained profile with that of the GTSE1 WT BAC.

### Protein purification

Full length GTSE1 was purified from insect cells using the protocol described in[23]. Briefly, insect cells expressing recombinant protein were harvested by centrifuging at 1800 rpm for 15 min and resuspended in 100mL ice-cold Buffer A (50 mM HEPES, pH 8.0, 300 mM NaCl, 5% glycerol supplemented with 2 mM TCEP and 1× protease inhibitor) followed by lysing the cells by sonication (Sonifier Cell Disruptor, Branson Ultrasonics Corp.) and then centrifuging at 30000 rpm for 30 min at 4°C. For the first step of affinity purification, the clarified supernatant was incubated with 1mL glutathione beads (Amintra) in a rolling shaker for 1 hour. The glutathione beads were then passed over gravity flow columns and washed in 3 iterations with 150 mL of Buffer A. An overnight incubation with GST Precission protease (purified in-house) in a volume of 5mL was performed to cleave off the GST tag. After eluting and concentrating the cleaved protein in Amicon Ultra-15 Centrifugal Filters 50K MWCO (Merck Millipore), size exclusion chromatography (SEC) was performed in a Superdex 200 10/300 column using gel filtration buffer (30 mM HEPES pH 8.0, 300 mM NaCl, 5% glycerol supplemented with 2 mM TCEP). The appropriate fractions were collected, concentrated to 5-10 μM and stored at −80°C.

For purification of GST-tagged EB1, *E. coli* Rosetta cells expressing GST-EB1 were grown to O.D._600_ of 0.7 and expression of recombinant protein was induced using 1 mM IPTG (Carl Roth) at 18°C overnight. Cells were harvested by centrifuging at 4000 rpm for 20 min and the pellet washed once with sterile PBS followed by resuspending and sonicating in 100 mL ice-cold GST binding buffer (25 mM HEPES pH 7.5, 300 mM NaCl, 1 mM EDTA, 5% glycerol) supplemented with 1% Triton X-100, 2 mM TCEP and protease inhibitor. The cell lysate was clarified by centrifuging at 30000 rpm for 30 min at 4°C. Affinity purification was carried out by passing the clarified lysate over a GSH column (GE Healthcare) using the ÄKTA Prime Plus system (GE Healthcare). After washing with GST binding buffer, the bound protein was eluted in GST binding buffer containing 20 mM Glutathione and concentrated to a volume of 2 mL in Falcon^BD^ concentrators with a 30000 MWDa cutoff at 3000 rpm. This was followed by performing SEC using GST binding buffer supplemented with 0.5 mM TCEP on a Superdex 200 10/300 column using an ÄKTA purifier (GE Healthcare). The fractions corresponding to the peak were pooled, concentrated and stored at −80°C.

His-tagged variants of GTSE1 (full length, 1-460 and 381-739) were used for *in vitro* phosphorylation reactions for identification of Cdk1 phosphosites on GTSE1. All expression constructs were cloned into *E. coli* Rosetta cells and purified as described in[54]. Briefly, bacteria expressing target protein were harvested and lysed by sonicating in ice-cold His binding buffer (20 mM Tris-HCl pH 8.0, 300 mM NaCl, 5% glycerol, 1 mM EDTA) supplemented with 10 mM Imidazole, 1% Triton X-100, 1 mM TCEP, DNAse and 1× protease inhibitor. The cell lysate was clarified by centrifugation at 30000 rpm and the clarified supernatant was passed through a gravimetric filter column to remove particulate matter, and then incubated for 3 hours with His beads (GE Healthcare) prewashed in His binding buffer. The beads were washed with 300 mL His washing buffer (20 mM Tris-HCl pH 8.0, 500 mM NaCl, 5% glycerol, 1 mM EDTA) supplemented with 30 mM Imidazole and 0.5 mM TCEP in three iterations, before being resuspended in His binding buffer, flash frozen and stored at −80°C.

### *In vitro* protein phosphorylation

For identification of phosphorylation sites on GTSE1, on-bead phosphorylation of full length GTSE1, 1-460 GTSE1 and 381-739 GTSE1 was performed using His-tagged protein purified from bacteria. Cdk1-cyclinB was a kind gift from Dr. Priyanka Singh and Dr. Andrea Musacchio (MPI Dortmund). Purified proteins were phosphorylated by Cdk1 at 4°C overnight, at a molar ratio of 1:100 (kinase:protein) in kinase buffer (20 mM HEPES pH 7.8, 150 mM NaCl, 5% glycerol supplemented with 2 mM MgCl_2_, 2 mM sodium orthovanadate and 10 mM ATP). Phosphorylation was confirmed by ProQ-Diamond^®^ staining of the protein. For using the phosphorylated protein for pulldown experiments, the kinase was first inactivated by adding 5 μM of Cdk1 inhibitor (RO-3306; Calbiochem) and 500 nM of Aurora A inhibitor (MLN8054; Sigma Aldrich) followed by incubation for 10 min on ice. In case of phosphorylation of EB1-GST using Cdk1, the kinase was removed after phosphorylated EB1-GST was bound to beads.

### GST Pulldowns

GSH beads (Amintra) were preblocked overnight in BSA blocking buffer (20 mM HEPES pH 7.5, 500 mM NaCl, 500 μg/mL BSA) as described earlier[77]. For the pulldown, 3 μM of the prey (phosphorylated or unphosphorylated protein) was mixed with 1 μM of the GST-tagged protein or the ‘bait’ and 5 μL of GSH beads in a total volume of 25 μL of the kinase buffer (20 mM HEPES pH 7.8, 150 mM NaCl, 5% glycerol). A sample (25%) was taken out as input before incubating the protein mixture on ice for 1 hour for the pulldown. The beads were then washed thrice with 250 μL of wash buffer (20 mM HEPES pH 7.0, 500 mM NaCl, 5% glycerol, 0.1% Triton X-100, 1 mM TCEP) and the protein was denatured by adding 20 μL 1× SDS-loading buffer before loading the samples on SDS gels. Pulldowns were quantified by calculating band intensities corrected for background using ImageJ.

### Cell culture and cell lines

All cell lines were grown in DMEM (PAN Biotech) containing 10% FBS (Gibco), 2mM L-glutamine (PAN Biotech), 100 U/mL penicillin, and 0.1 mg/ml streptomycin (PAN Biotech) at 37°C in 5% CO_2_. U2OS cells stably expressing RNAi-resistant wild-type GTSE1-LAP were described previously[53]. To obtain U2OS cells expressing the GTSE1-GFP or GTSE1-14A-GFP transgenes, the corresponding BACs were transfected into U2OS cells using the Effectene kit (Qiagen) following the manufacturer’s instruction. Clonal lines expressing the BAC transgene were selected with Blasticidin (Sigma Aldrich). The GTSE1^WT^ and GTSE1^14A^ clones expressing the GTSE1-GFP and GTSE1-14A-GFP BAC transgenes close to endogenous level, respectively, were selected for phenotypic analysis. Transient transfections of GTSE1 point mutants were performed in ibidi chambers using 750 ng DNA and 0.75 μl Lipofectamine 2000 (Invitrogen) according to the manufacturer’s protocol.

### Antibodies

Antibodies used: goat anti-GFP (MPI Dresden; described in[78]), rabbit anti-GTSE1 (custom generated; described in [53]), mouse anti-α-tubulin (DM1α, Sigma-Aldrich), mouse anti-CHC (X22, ab2731, Abcam), rat anti-EB1 (KT-51, Absea Biotechnology), mouse anti-MCAK/Kif2C (1G2, Abnova Corp.), human nuclear antibodies to nuclear antigens centromere autoantibody (CREST) (CS1058, Europa Bioproducts Ltd), rabbit anti-CEP135 (MPI Dresden, described in[79]), donkey anti-human antibodies conjugated to Cy5 or Texas red (Jackson ImmunoResearch laboratories; #709-175-149 and #709-075-149), donkey anti-rat Alexa488 (Bethyl; A110-337D2, donkey anti-rat Alexa594 (Bethyl; A110-337D4), donkey anti-rabbit Alexa488 (Thermofisher; A-21206), donkey anti-rabbit Alexa594 (Bethyl; A120-208D4), donkey anti-rabbit Alexa650 (Bethyl; A120-208D5), donkey anti-mouse Alexa488 (Bethyl; A90-337D2), donkey anti-mouse Alexa594 (Bethyl; A90-337D4), donkey anti-mouse Alexa647 (Invitrogen; A31571), donkey anti-goat Alexa488 (Jackson Immunoresearch; 705 545 147), donkey anti-goat HRP (Santa Cruz; SC-2020), sheep anti-mouse HRP (Amersham; NXA931-1ml), donkey anti-rat (Amersham; check make) donkey anti-rabbit HRP (Amersham; NXA934-1ml).

### Immunoprecipitation (IPs)

Cells at ^~^70% confluency were arrested in prometaphase using 10 μM S-trityl-L-Cysteine (STLC; Sigma Aldrich) for 16 hours and collected using mitotic shake off. They were then pelleted down by centrifuging at 1000 rpm for 5 min and then washed with PBS supplemented with 10 μM MG-132 (Calbiochem). After centrifugation, the cells were resuspended in complete DMEM medium supplemented with 10 μM MG-132 and incubated in an incubator maintained at 37°C and 5% CO_2_ for 90 min to allow the cells to reach metaphase. Following this, the cells were once again washed, lysed in cell lysis buffer (50 mM Na_2_HPO_4_, 150 mM NaCl, 10% glycerol, 1% Triton X-100, 1 mM EGTA, 1.5 mM MgCl_2_, 50 mM HEPES pH 7.2, 1 mM DTE) supplemented with 2× protease inhibitor mix (SERVA), and PhosStop (EASY Pack, Roche)) and incubated on ice for 15 min. The cell suspension was then clarified by centrifuging at 13000 rpm at 4°C for 15 min. Total protein concentration of the cell extract was calculated using Bradford reagent. For performing immunoprecipitation, 1 mg total protein in a volume of 1 mL was used. A 4% sample was taken out as input. To the remaining cell lysate, 1-2 μg of indicated antibodies were added and the mixture incubated at 4°C for 90 min. Subsequently, pre-washed dynabeads coupled to protein G (Invitrogen) were added to the lysate and the mixture incubated at 4°C for 4 hours. The beads were then washed thrice with 1 mL of the appropriate lysis buffer, resuspended in SDS-loading buffer and boiled at 95°C for 5 min before analyzing by SDS-PAGE and western blotting.

### RNAi

siRNA against hGTSE1 (5’-GAU UCA UAC AGG AGU CAAA-3’), EB1 (5’-UUC GUU CAG UGG UUC AAGA-3’), and control siRNA (Silencer negative control 2; AM4637) were purchased from Ambion. Approximately 35,000 U2OS cells were added to prewarmed media in 24-well plates or 8-well imaging chambers (ibidi), and transfection complexes containing 2.5 μl Oligofectamine and siRNA were added immediately afterward. Media was changed after 6–8 hours. Final concentrations of 80 nM and 100 nM RNAi were used for GTSE1 and EB1, respectively.

For depleting endogenous GTSE1, EB1 in U2OS cells, reverse transfection using 80 nM of silencer RNA (siRNA) for GTSE1 and 100 nM of siRNA for EB1 was performed using Oligofectamine (Invitrogen). Briefly, transfection complexes consisting of siRNA and Oligofectamine were prepared by mixing 2.5 μL Oligofectamine with siRNA in OptiMEM (Invitrogen) at room temperature for 20 min. This was added to roughly 35000-40000 cells in prewarmed medium in 24-well plates (Sarstedt) containing glass coverslips, or 8-well imaging chambers (ibidi), and the volume made up to 500 μL. The cells were left in the humidified incubator at 37°C for 8 hours after which the media was replaced. All experiments were performed 48 hours after transfection.

Clathrin depletion in human cells was performed using a forward transfection approach. Cells were seeded into a 3.5 cm dish with coverslips and grown until 75% confluent. Prior to transfection, medium was exchanged for 1.8 mL OptiMEM supplemented with 100 U/mL penicillin and 0.1 mg/mL streptomycin (PAN Biotech, Pen/Strep mix). A 200 μL transfection mix containing 3 μL Lipofectamine RNAi Max (Invitrogen) and the siRNA was prepared in OptiMEM (Invitrogen), and incubated for 25 min at RT before addition to the cells. Experiments were performed 66 hours after transfection. Clathrin heavy chain (CHC) siRNA was used at 50 nM final concentration. When necessary, cells transfected with Control or CHC siRNA for 66 hours using the forward transfection approach were submitted to a second round of reverse transfection with Control, GTSE1 (100 nM), or CHC (100 nM) siRNAs for an extra 48 hours.

### Western blot

For western blotting after RNAi, cells were harvested by directly adding hot Lamelli buffer in 24-well plates or ibidi chambers as required. The proteins were separated on SDS-PAGE gels and transferred onto nitrocellulose membranes. Membranes were blocked using either 5% milk prepared in PBS containing 0.1 % Tween (PBS-T) or in the case of CHC, using 5% bovine serum albumin (BSA) in PBS-T for 1 hour at room temperature. The membrane was incubated with the indicated primary antibodies, followed by secondary antibodies coupled to horseradish peroxidase. Protein signal was detected using chemiluminescence with the ECL Prime Western Blotting Detection Reagent™ (GE Healthcare) according to manufacturer’s instructions. After incubation with the ECL blotting reagent, images were acquired using the ChemiDoc™ MP Imaging System (BioRad).

### Mass spectrometry

For MS analysis, IPs and *in vitro* phosphorylation reactions were performed in triplicates to obtain reliable label-free data. Liquid chromatography coupled with mass spectrometry was used to assess the phosphorylation status of GTSE1. *In vitro* phosphorylated GTSE1 (full-length, 1-460 or 381-739) using Cdk1 as kinase and its unphosphorylated controls were reduced, alkylated and digested with LysC/Trypsin and prepared for mass spectrometry as previously described[80]. Obtained peptides were separated on a Thermo Scientific™ EASY-nLC 1000 HPLC system (Thermo Fisher Scientific™, Odense, Denmark) using a 45 min gradient from 5-60% acetonitrile with 0.1% formic acid and directly sprayed via a nano-electrospray source in a quadrupole Orbitrap mass spectrometer (Q ExactiveTM, Thermo Fisher Scientific™) [81]. The Q Exactive™ was operated in data-dependent mode acquiring one survey scan and subsequently ten MS/MS scans[82]. Resulting raw files were processed with the MaxQuant software (version 1.5.2.18) using GTSE1 and Cdk1 for the database search. Deamidation (NQ), oxidation (M) and phosphorylation (STY) were given as variable modifications and carbamidomethylation (C) as fixed modification [83]. A false discovery rate cut off of 1% was applied at the peptide and protein levels and as well on the site decoy fraction [83]. Identified phospho peptides were further separated in class I sites where the calculated localization probability on a specific residue was >75% and class II sites with localization probabilities <75%. Immunopreciptiated GTSE1 was directly digested on beads [60] and peptides subsequently treated as described above using a 90 min gradient (5-60% acetonitrile with 0.1% formic acid) instead of 45 min.

### Immunofluorescence

Cells grown on coverslips were fixed using pre-cooled methanol (Sigma Aldrich) for 10 min at −20°C, followed by washing and rehydration in PBS for 10 min. Next, the cells were blocked for 1 hour in 5% BSA prepared in PBS followed by incubation for 1 hour in the dark with appropriate dilutions of primary antibodies (prepared in 5% BSA) in a humidified chamber at 37°C. After washing the coverslips thrice in PBS to remove excess primary antibody, they were similarly incubated with corresponding secondary antibodies, and washed thrice to remove excess antibody. Coverslips were mounted on glass slides using Prolong Gold containing DAPI (Invitrogen).

Images used for quantifying astral microtubule lengths in metaphase were acquired using the 60×/1.4 NA Plan-Apochromat Oil Objective (Zeiss) on the DeltaVision Imaging System (GE Healthcare) equipped with an sCMOS camera (PCO edge 5.5). Serial Z-stacks of 0.25 μm thickness were acquired and then deconvolved using SoftWoRx 6.1.1 software. All other images for quantifying inner spindle intensity, microtubule lengths in prometaphase and spindle tilt were taken using the 3i Marianas™ spinning disk confocal system (Intelligent Imaging Innovations Inc.) equipped with Axio Observer Z1 microscope (Zeiss), Plan-Apochromat 60×/1.4 NA Oil Objective, M27 with DIC III Prism (Zeiss), Orca Flash 4.0 sCMOS Camera (Hamamatsu Photonics) and controlled by Slidebook Software 6.0 (Intelligent Imaging Innovations Inc.).

### Live cell imaging

For identifying kinase dependency on phosphorylation of GTSE1 in mitosis, GTSE1^WT^ cells were treated with appropriate dilutions of specific kinase inhibitors in CO_2_-independent visualization medium (Gibco) supplemented with 10% FBS, 2mM L-glutamine and penicillin/streptomycin as mentioned earlier. Aurora A inhibitor (MLN8054; Sigma Aldrich) and Aurora B inhibitor (ZM447439; EMD Millipore Corp.) were added at 500 nM and 2 μM respectively for 90 min while 100 nM of Plk1 inhibitor (BI 2536; Enzo Life Sciences) was added for one hour prior to imaging. For inhibiting Cdk1, 9 μM RO-3306 (Calbiochem) was used. Cells were imaged in prometaphase first for 20 seconds followed by replacing the media with fresh visualization medium containing RO-3306 and imaging for 180 seconds. A z-stack consisting of 3 sections at intervals of 0.4 μm was acquired every 1 second. Cells transfected with phosphonull GTSE1 mutants were imaged using similar parameters.

To visualize DNA for measuring the time from NEBD to anaphase, 1 hour prior to imaging, 500 nM of the cell permeable DNA dye called siR-DNA (Spirochrome) and 1 μM Verapamil, a sodium channel efflux inhibitor, were added to the visualization media. Serial z-stacks of 2 μm were acquired every 2 min for 12 hours in a heated chamber (37°C). For visualizing microtubules for analyzing spindle positioning defects, 1 hour prior to imaging, 125 nM of siR-tubulin (Spirochrome) and 1 μM Verapamil were added to the visualization media. All live cell imaging was performed on the DeltaVision Imaging System (GE Healthcare) using the 40×/1.3 NA UPLFLN Oil Objective.

For studying microtubule dynamics in mitosis, GTSE1^WT^ and GTSE1^14A^ cells dually expressing EB3-mCherry were used and imaged using the 60× Oil Objective on the 3i Marianas™ spinning disk confocal system. Three serial z-stacks of 0.4 μm were acquired every second for two min at 37°C.

### Image analysis and quantification

Image quantification was performed on unmodified 16-bit z-series images using ImageJ and Imaris 7.6.4 32-bit software (Bitplane). On Imaris, all images were opened and analyzed in the ‘Surpass’ mode in the software.

For determining colocalization between GTSE1 and EB3, the ‘Spots’ function with a diameter threshold of 0.5 μm was applied to the EB3 channel to identify comets, and the intensities of EB3 and GTSE1 at identified comets was extracted. An average value of background intensity for both channels was obtained by placing three points on the microtubule lattice (identified from faint EB3 signal). Background subtracted intensities of EB3 were thresholded based on a value determined by visually confirming correct annotation of ‘faint’ EB3 comets. GTSE1 intensities at corresponding spots was calculated in the same manner, without thresholding. For colocalization analysis during mitosis, a region of interest lying outside the spindle, over astral microtubules was chosen.

For making line scans to compare GTSE1 and EB3 localization at microtubule plus ends across the cell cycle, maximum projection images of cells expressing GTSE1-GFP and EB3-mCherry were opened on ImageJ, and ^~^4 μm long lines were manually drawn along EB3 labelled microtubule ends. All lines began on the microtubule lattice and terminated outside the comet. The absolute intensities of EB3 and GTSE1 were extracted using the Plot Profile plugin of ImageJ. All line scan intensity profiles were aligned to the position of the peak intensity of the EB3 channel and normalized before plotting using MATLAB scripts.

To calculate the number of GTSE1 comets after transient transfection, stills from live-cell imaging of equally transfected cells were opened on Imaris. A ROI of fixed dimensions was placed over astral microtubules. The ‘Spots’ function with a diameter threshold of 0.5 μm was used to identify GTSE1 comets. The average background GFP intensity was calculated from three randomly placed points in the cytoplasm. Background corrected intensities were thresholded using a value 1.5× greater than the average background.

Raw images of cells stained for EB1, tubulin and CEP-135 were used for calculating microtubule lengths by adapting the protocol outlined in (Stout et al., 2011) and writing MATLAB functions and scripts. The 3D position of the spindle poles (*P_x_, P_y_, P_z_*) was determined by using the ‘Measurement’ function on the CEP-135 signal. The ‘Spots’ function was used to determine spherical coordinates of all EB1 comets (*C_x_, C_y_, C_z_*) using the spot diameter thresholded at 0.5 μm with application of occasional manual filtering to eliminate spots not also visibly associated with tubulin. These coordinates were extracted using the ‘Statistics’ function on Imaris. MATLAB scripts were used to calculate distances between all comets to both spindle poles using the formula 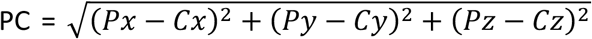. The shorter of the two distances was used as a proxy for microtubule length. The spindle length ‘*l*’ was determined by calculating the distance between the two poles according to the equation written above. The spindle width ‘*w*’ was defined as the distance between the outermost kinetochore fibers on opposite ends of the spindle attached to DNA and was manually determined using the tubulin channel. This was used to determine the spindle angle θ, defined as tan^−1^(*w/l*). Next, the angle φ between the line formed by a comet and its closest pole, and the line formed by the two spindle poles, was calculated using the cosine law. An EB1 comet was associated with an astral microtubule if ϕ was greater than θ. The mean length of astral microtubules per cell and the total number of astral microtubules per cell were calculated. For cells in prometaphase, owing to cytoskeleton and spindle morphology, there is no defined spindle angle. Therefore, the lengths of all microtubules were calculated as described above. All analyses were performed in Microsoft Excel and MATLAB R2019a.

For calculating the intensity of tubulin in the inner spindle, the spindle was detected using the ‘Surface’ module on Imaris using a surface detail value of 1.0 on the tubulin channel. The total fluorescence intensity of tubulin in a manually placed ROI over the spindle was obtained through the ‘Statistics’ option. The volume corrected background intensity was calculated by multiplying the spindle volume with the average tubulin intensity of four manually placed spots in the cytoplasm that lacked polymerized tubulin. The tubulin intensity in the inner spindle corrected for the background was calculated.

For semi-automatic detection of number of microtubules reaching the cortex, maximum projections of 16-bit raw images were used to determine the outline of cells using the cytoplasmic intensity of the CEP-135 fluorescence. A line of 1.05 μm thickness was drawn towards the inside of the cell boundary and the corresponding intensities of tubulin across that line were extracted using the Plot Profile plugin on ImageJ. A background value obtained by averaging the intensities at 3 random points lacking tubulin along the cortex-adjacent line was subtracted from absolute intensities at all points. Background corrected tubulin intensites were subjected to a threshold value, determined from visual inspection of correctly annotated microtubules in at least eight cells per experiment. The number of microtubules thus obtained was divided by the length of the cortex of the corresponding cell.

For calculating the angle α made between the spindle and the long cell axis at different time points, unmodified 16-bit videos were opened on ImageJ and the long cell axis was defined 8 min prior to NEBD. The angle α was measured using the Angle tool of ImageJ software.

### Microtubule dynamics

Microtubule dynamics were analyzed from EB3-mCherry movies using the software plusTipTracker that has been made publicly available by the Danuser laboratory [84]. The same parameters were used for all movies: maximum gap length, fifteen frames; minimum track length, three frames; search radius range, 5-25 pixels; maximum forward angle, 60° maximum reverse angle, 15° maximum shrinkage factor, 1.0; fluctuation radius, 1.5 pixels; and time-interval 1 s. Data collected from the analysis included mean microtubule growth velocity, mean microtubule growth-track lifetime, number of bgaps and the growth time before bgap. Catastrophe frequency was calculated as the number of bgaps (corresponding to shrinkage events) divided by the growth time before bgap.

### Statistical analysis

Statistical significances were determined by performing two-tailed Student’s t tests, and Welch’s correction was applied if the standard deviation of the distributions under comparison were more than 2× different from each other. Significances were calculated using Microsoft Excel, Graphpad Prism 7 and MATLAB 2019a.

**Supplementary Figure 1:**
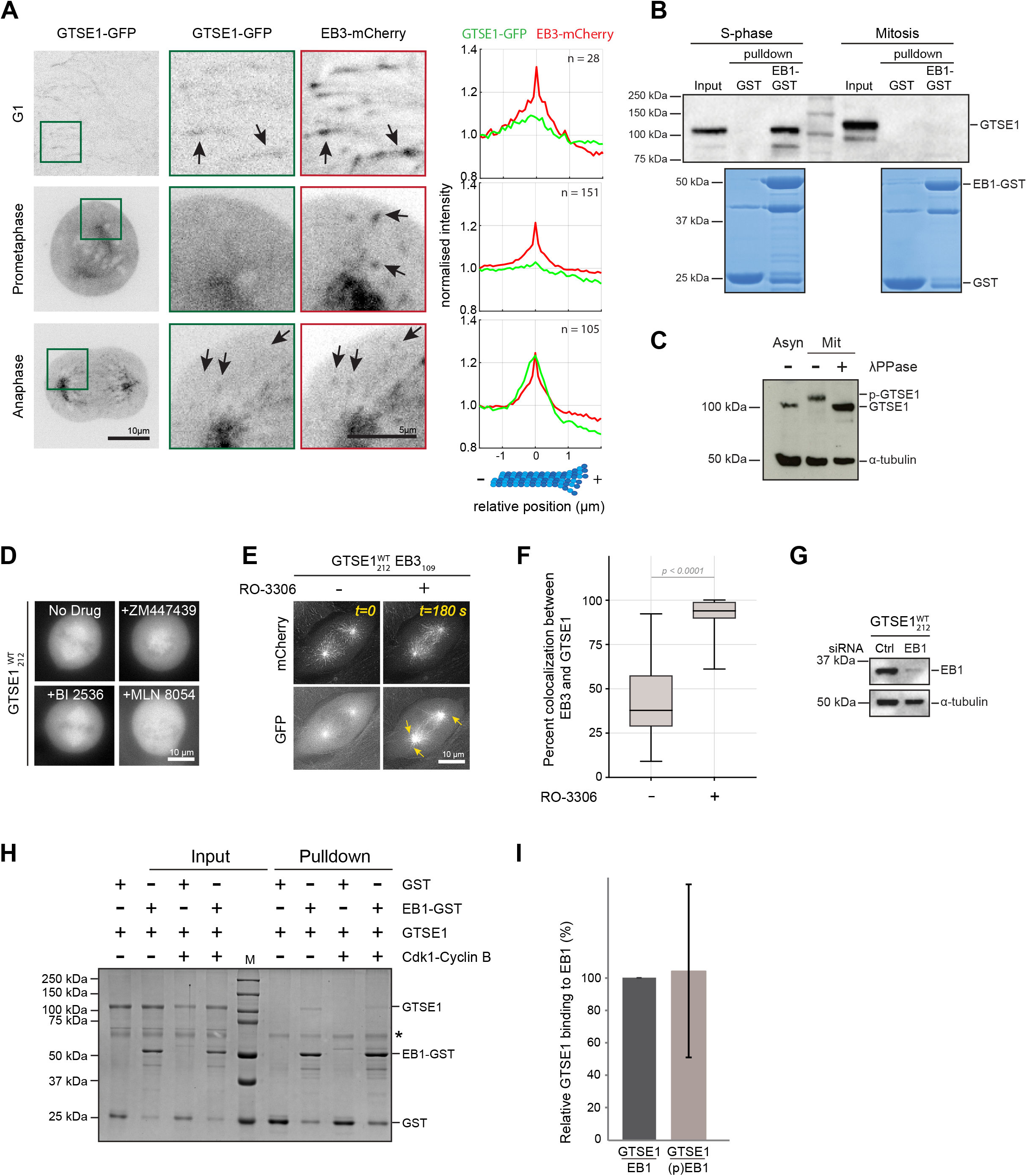
Cdk1 removes GTSE1 from growing microtubule plus-ends in mitosis. **(A)** Live cell images of U2OS cells stably expressing GTSE1-GFP from a BAC (green) and EB3-mCherry from a plasmid (red) in different cell cycle stages. Scale bar 10 μm. Magnified insets show GTSE1 (middle) and EB3 (right) with arrows indicating comets, scale bar 5 μm. Averaged line scan intensity profiles of microtubule plus-ends show relative intensities of GTSE1-GFP (green) and EB3-mCherry (red), centered at the maximum EB3 intensity. Between 28-151 microtubule plus ends from 2 cells in 2 experiments expressing comparable levels of both transgenes were analyzed. **(B)** Immunoblot using anti-GTSE1 antibody of EB1-GST pulldown using lysates from U2OS cells either arrested in S-phase using Thymidine or in prometaphase using STLC. Shown below is Coomassie Blue staining of GST and EB1-GST pulled down in both phases. **(C)** GTSE1 is hyperphosphorylated in mitosis. Immunoblot of U2OS lysates from asynchronous and mitotic cells treated (or not) with λ-phosphatase (λ-PPase). Immunoblot using anti-GTSE1 and anti-tubulin antibodies. **(D)** Stills from live cell imaging in mitosis of U2OS cells expressing GTSE1-GFP from a BAC (GTSE1^WT^_212_), treated with inhibitors of mitotic kinases Aurora B (ZM447439), Plk1 (BI 2536) and Aurora A (MLN 8054) and MG-132 to prevent exit from mitosis. Scale bar 10 μm. **(E)** Single frames from live cell microscopy of U2OS cells dually expressing GTSE1-GFP from a BAC and EB3-mCherry from a plasmid (GTSE1^WT^_212_EB3_109_), before and 3 minutes after treatment with a Cdk1 inhibitor (RO-3306). EB3 is shown on the top and corresponding GTSE1 signal is shown at the bottom. Arrows indicate comets. Scale bar 10 μm. **(F)** Box plot showing percent colocalization between EB3 and GTSE1 before and after addition of RO-3306 from experiment shown in (B). n = 21 cells per condition over 3 experiments. P-value from unpaired t-test with Welch’s correction. Error bars indicate s.e.m. **(G)** Western blot of GTSE1^WT^_212_ cells treated with control (Ctrl) or EB1 siRNA for 48 hours. Immunoblot using anti-GTSE1 and anti-tubulin antibodies. **(H)** Coomassie Blue stained gel of a GST pulldown assay using EB1-GST phosphorylated (or not) with Cdk1-Cyclin B as bait to pull down purified GTSE1. ‘*’, GSH beads blocked using BSA. **(I)** Quantification of relative GTSE1 binding to EB1 before and after phosphorylation of EB1 by Cdk1. n = 3 experiments. Error bars represent s.d.

**Supplementary Figure 2:**
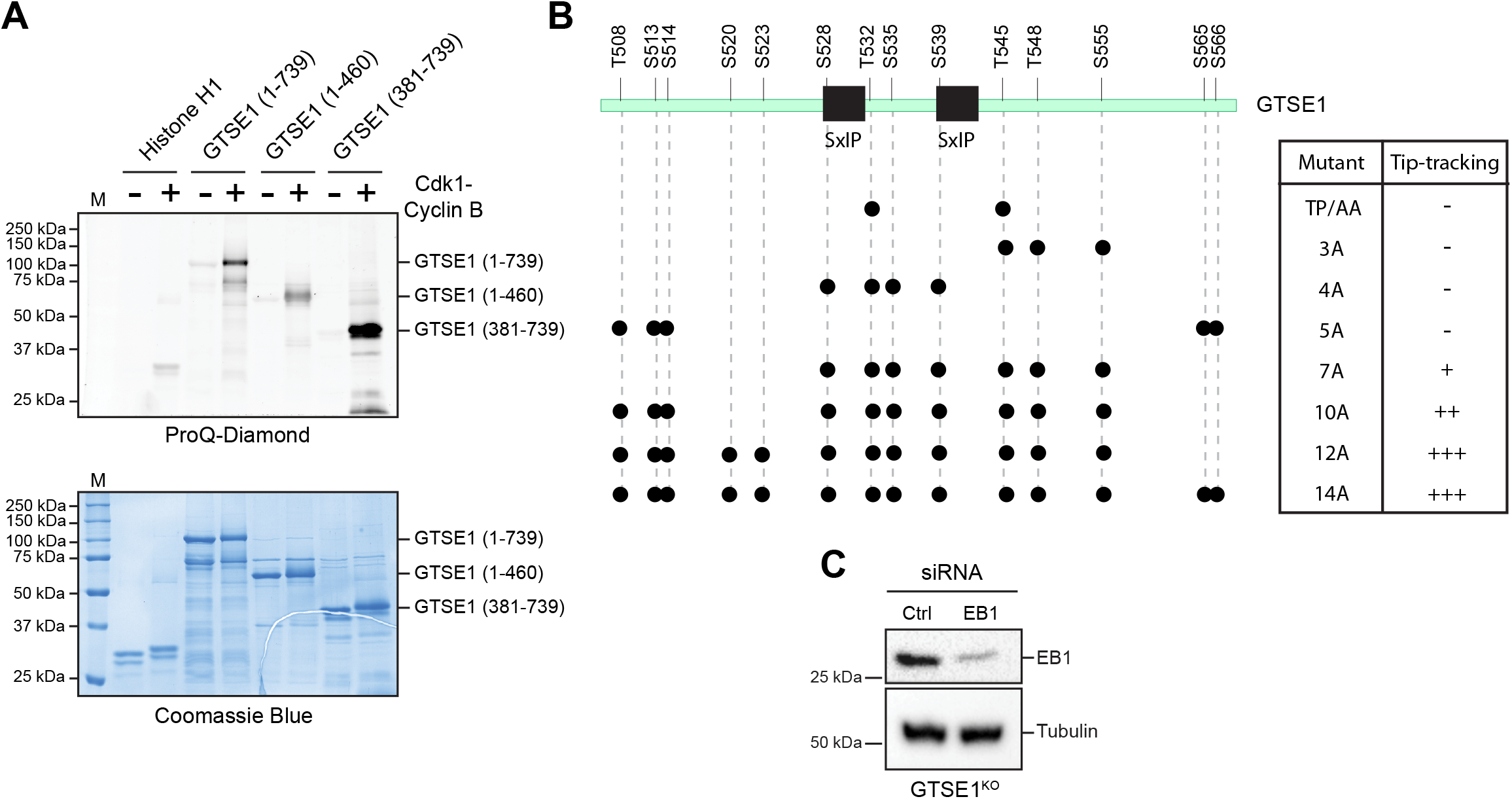
Phosphosite mutations in GTSE1 and impact on plus-end tracking. **(A)** Cdk1-cyclinB phosphorylation of His-tagged GTSE1 and its fragments. Histone H1 was used as positive control for Cdk1 activity. The upper gel shows phosphorylated proteins detected by ProQ-Diamond staining. The same gel was also stained with Coomassie Blue to detect loading of proteins. **(B)** Schematic of GTSE1 with residues targeted for mutation in this study. The table shows the (Ser/Thr)Ala mutants of GTSE1 generated and their corresponding plus-end tracking behaviour in mitosis as observed by live imaging after transient transfection in GTSE1^KO^ cells. **(C)** Western blot of GTSE1^KO^ cells treated with control (Ctrl) or EB1 siRNA for 48 hours. Immunoblot using anti-GTSE1 and anti-tubulin antibodies.

**Supplementary Figure 3:**
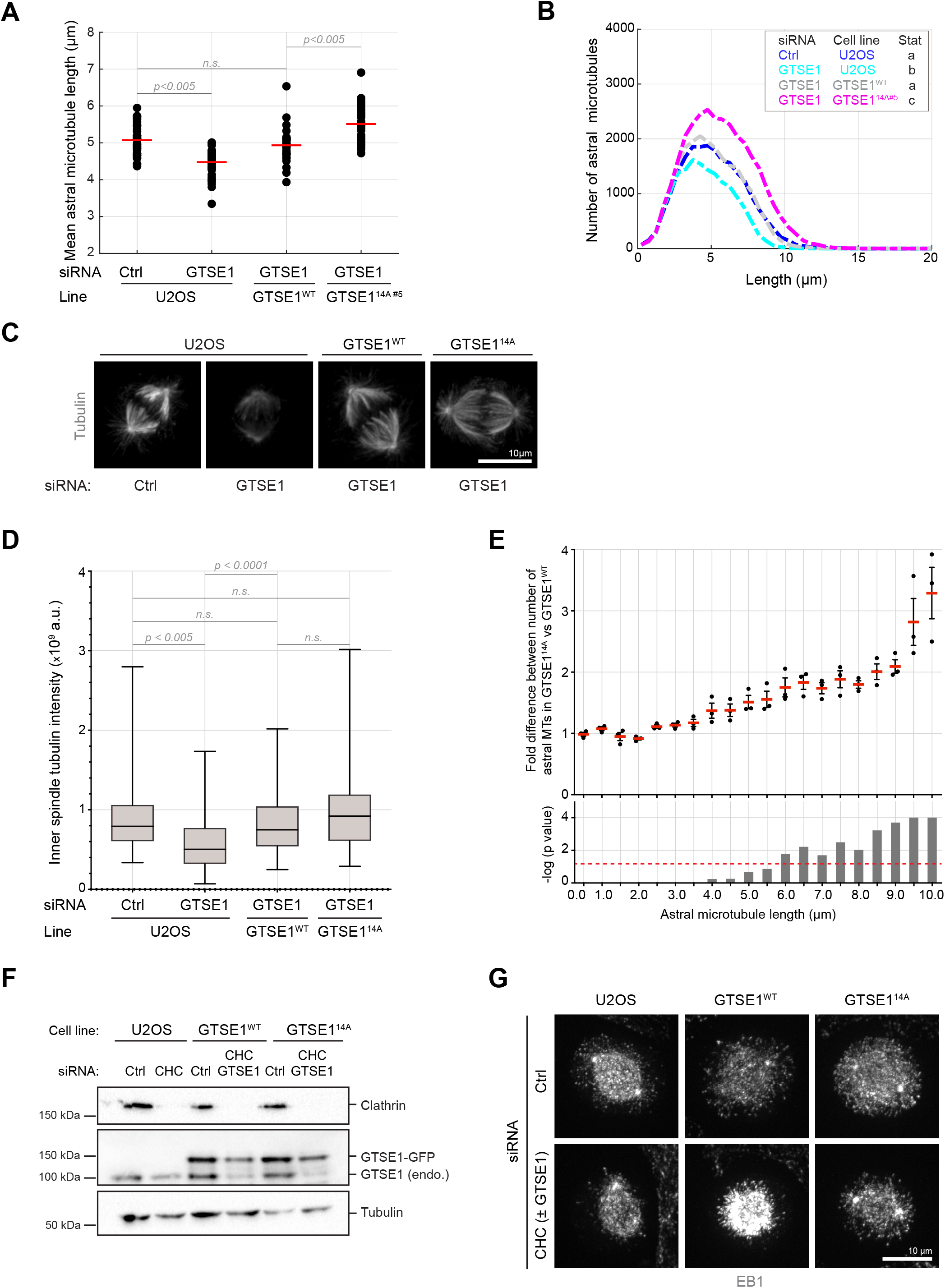
Cdk1-dependent removal of GTSE1 from growing microtubule plus-ends stabilizes long astral microtubules, but not inner spindle microtubules, in metaphase. **(A)** Dot plot showing mean astral microtubule length in U2OS, GTSE1^WT^ and a second independent clone expressing GTSE1-14A (GTSE1^14A^). Red line indicates mean. n = 37 cells from 3 independent experiments, pooled for representation. P-values from unpaired two-tailed student’s t-test. **(B)** Distributions of number of astral microtubules as a function of their lengths, for the experiment shown in (A). n = 37 cells from 3 independent experiments, pooled for representation. Inset shows the significances (different letters indicate groups differing significantly). **(C)** Immunofluorescence images of metaphase U2OS, GTSE1^WT^ and GTSE1^14A^ after control (Ctrl) or GTSE1 RNAi, stained with anti-tubulin antibody. Scale bar 10 μm. **(D)** Box plot showing intensity of tubulin in the inner spindle from experiment shown in (E). n = 55-76 cells, from 3 independent experiments. P-values from unpaired t-test with Welch’s correction. **(E)** Quantification of the fold difference in the number of astral microtubules of different lengths between metaphase in GTSE1^WT^ and GTSE1^14A^. P-values from one-way ANOVA. The horizontal red dashed line indicates a p-value of 0.05. **(F)** Western blot of cell lysates of U2OS, GTSE1^WT^ and GTSE1^14A^ treated with control (Ctrl), clathrin heavy chain (CHC) alone or CHC in combination with GTSE1 siRNAs. Immunoblots using anti-CHC, anti-GTSE1 and anti-tubulin antibodies. **(G)** Immunofluorescence images in metaphase of U2OS, GTSE1^WT^ and GTSE1^14A^ cells treated with control (Ctrl), CHC siRNA alone (U2OS) or a combination of CHC and GTSE1 siRNA (in GTSE1^WT^ and GTSE1^14A^). Cells were fixed and stained with antibodies against EB1. Scale bar 10 μm.

**Supplementary Figure 4:**
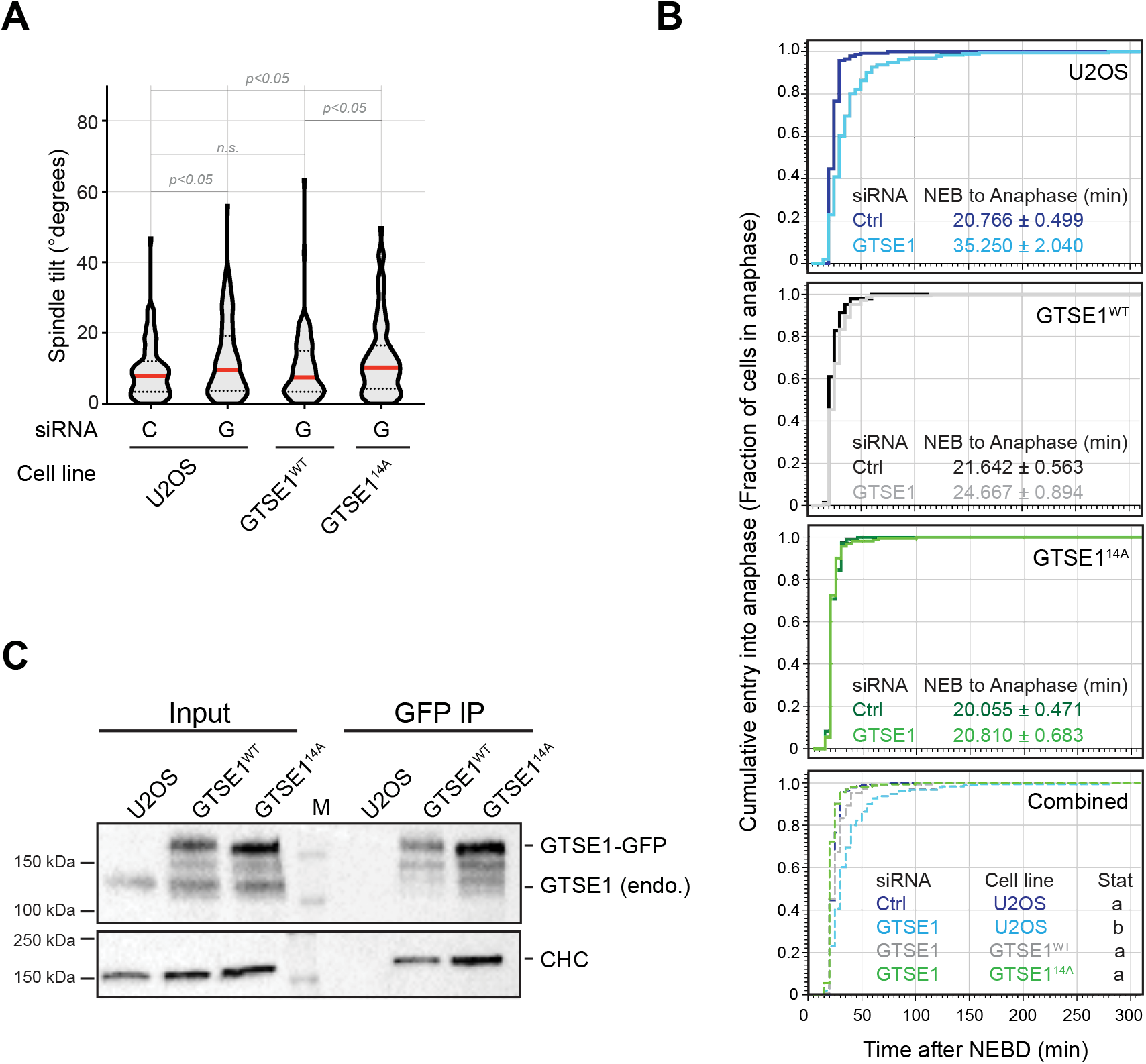
Hyperstabilization of astral microtubules does not affect progression to anaphase. **(A)** GTSE1^14A^ cells are defective for metaphase spindle alignment. Violin plot with quantification of the angle made by the spindle with the substrate (spindle tilt) in U2OS, GTSE1^WT^ and GTSE1^14A^ cells, transfected with control (C) or GTSE1 (G) siRNA. n > 99 cells from 3 experiments per condition. Red line indicates the median. P-values calculated from unpaired t-test with Welch’s correction. **(B)** Fraction of cells that have entered anaphase as a function of the time (min) after NEBD. The mean (± s.e.m) time between NEBD and anaphase onset is indicated in the insets. U2OS, GTSE1^WT^ and GTSE1^14A^ cells were transfected with control (Ctrl) or GTSE1 siRNA. n = 109-191 cells per condition over 3 independent experiments. Data was pooled for representation. Inset in ‘Combined’ shows the significances (different letters indicate groups differing significantly). P-values from unpaired t-test. **(C)** Immunoblots of cell lysates (input) and immunoprecipitations (IP) of GFP in U2OS, GTSE1^WT^ and GTSE1^14A^ cells. Immunoblots with anti-GTSE1 and anti-CHC antibody.

**Video 1: Inhibition of Cdk1 restores EB1-dependent GTSE1 plus-end tracking in mitosis.** U2OS cells expressing GTSE1-GFP as a transgene from a BAC were treated with Control (Ctrl) siRNA and imaged at 1 second-intervals for 20 seconds before adding small molecule inhibitor against Cdk1 (RO-3306) and imaging for 3 min. Video shown at nine frames per second.

**Video 2: Inhibition of Cdk1 restores EB1-dependent GTSE1 plus-end tracking in mitosis.** U2OS cells expressing GTSE1-GFP as a transgene from a BAC were treated with EB1 siRNA and imaged at 1 second-intervals for 20 seconds before adding small molecule inhibitor against Cdk1 (RO-3306) and imaging for 3 min. Video shown at nine frames per second.

**Video 3: Cdk1 inhibition in mitosis reversibly restores GTSE1 plus end-tracking in mitosis.** U2OS cells expressing GTSE1-GFP as a transgene from a BAC were imaged as above at 1 second-intervals for 10 seconds before adding RO-3306 and imaging for 25 seconds. The drug was washed out and cells imaged again for 50 seconds. Video shown at six frames per second.

**Video 4: A phospho-mutant of GTSE1 displays plus-end tracking in mitosis.** U2OS cells knocked out of GTSE1 (GTSE1^KO^) were transiently transfected with plasmids expressing wildtype (WT) GTSE1-GFP or GTSE1-14A-GFP, and imaged at 1 second-intervals for 20 seconds. Video shown at nine frames per second.

**Video 5: GTSE1-14A-GFP plus-end tracks in mitosis in an EB1-dependent manner.** U2OS cells knocked out of GTSE1 (GTSE1^KO^) were treated with control (Ctrl) or EB1 siRNA and transiently transfected with the plasmid expressing GTSE1-14A-GFP. Cells in mitosis were imaged at 1 second-intervals for 20 seconds and displayed at nine frames per second.

**Video 6: Cdk1 inhibition in mitosis reversibly restores GTSE1 plus end-tracking in mitosis.** U2OS clones stably expressing wildtype (WT) GTSE1-GFP (GTSE1^WT^) or GTSE1-14A-GFP (GTSE1^14A^) were imaged in interphase and mitosis at 1 second-intervals. Video is displayed at 9 frames per second.

